# A semi-automated pipeline for quantitation of Pax7+, myonuclei, and cross-sectional area by fiber type

**DOI:** 10.64898/2026.06.03.729866

**Authors:** Helia G. Megowan, Madeline Luu, Adam Shuaib, Kaitlyn B. Augienello, Adam C. Fries, Jake Searcy, Hans C. Dreyer

## Abstract

Manual analysis of skeletal muscle cross-sections is time-consuming and subject to error and user bias. To overcome these limitations, we developed and validated a semi-automated, quantitative, and reproducible image-analysis pipeline specifically tailored to quantify Pax7+ satellite cells, myonuclei, and cross-sectional area by fiber type. The workflow combines FIJI/ImageJ-based image preprocessing with CellProfiler, Cellpose, and a custom Python script to process and analyze immunohistological images of muscle tissue cross-sections. Outcomes include Pax7+ satellite cells and myonuclei quantified per fiber by fiber type, along with cross-sectional area, perimeter, and fiber type classification. This semi-automated approach provides a robust and efficient platform for high-throughput analysis of muscle tissue cross-sections from large datasets.

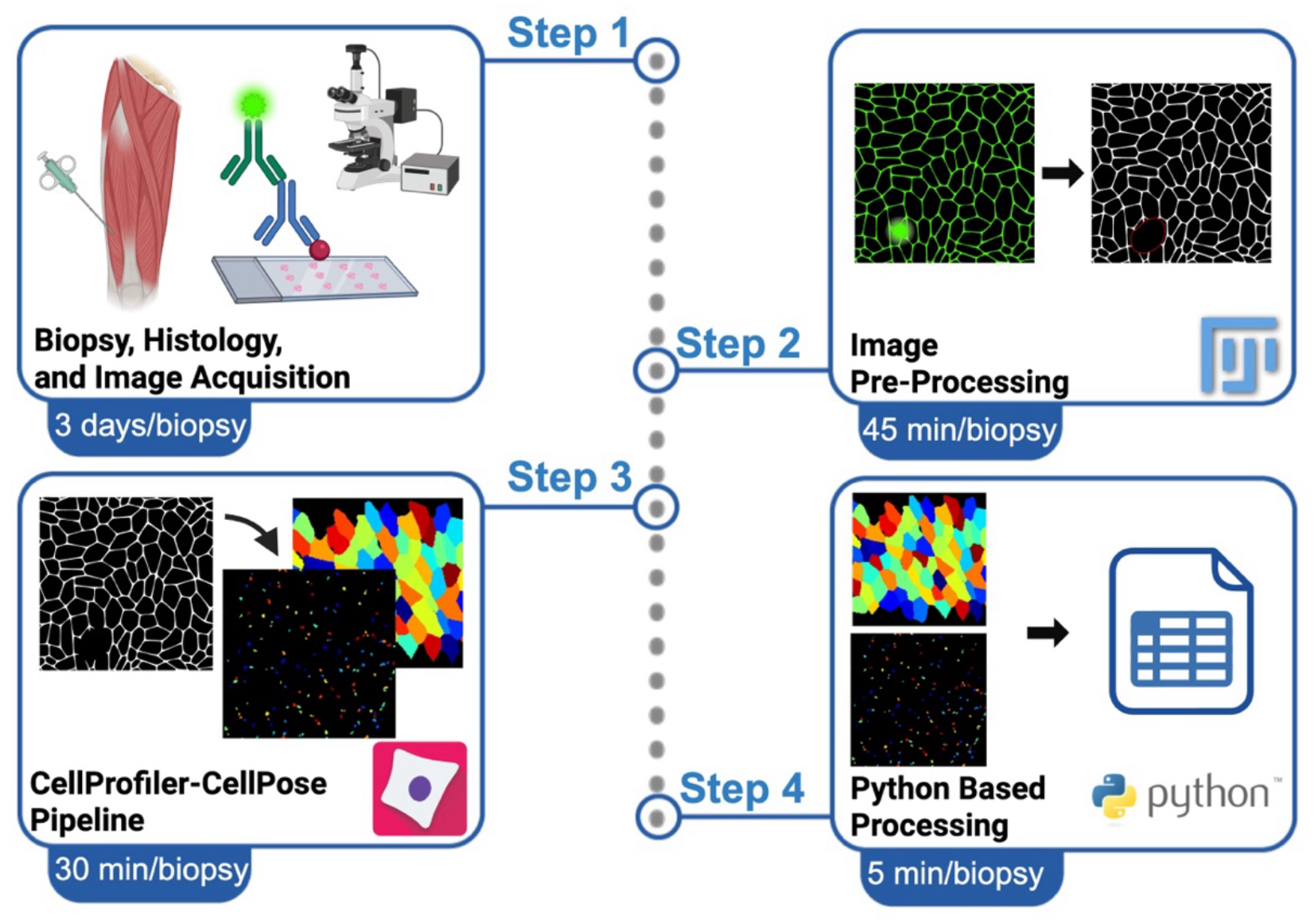

## Introduction

Histological evaluation of skeletal muscle reveals how tissue structure relates to baseline physiology and experimental responses^1^. Immunohistochemistry, using labeled antibodies to detect specific proteins, combined with high-resolution imaging, maps the cellular and subcellular organization of muscle. Muscle fibers are highly plastic—showing altered cross-sectional area (CSA), shifts in myonuclear number and position, and modulation of Pax7+ satellite cell pools—in response to internal and external cues. These histological endpoints are routinely used as structural surrogates for muscle functional capacity and adaptative potential.

Precise quantification of Pax7+ satellite cells is essential for understanding of skeletal muscle adaptations, including myonuclear accretion. Satellite cells give rise to new myonuclei, making them relevant to the ongoing assessment of the myonuclear domain hypothesis, which posits that each myonucleus regulates a defined cytoplasmic volume^2–4^. Emerging data indicate the domain may align more closely with fiber perimeter than with cross-sectional area, supporting use of perimeter as an outcome.^5^ Retention of newly accreted myonuclei is implicated in “muscle memory,” whereby previously trained muscle regains size and strength more rapidly than naïve muscle^6,7^. Together, these measures provide mechanistic insight into muscles’ responses to interventions, providing foundational evidence for their use in clinical applications^8^.

A common approach quantifies muscle structural changes on cross-sections cut perpendicular to fiber long axes, which samples many individual fibers (each fiber is a single muscle cell) and therefore better represents the muscle. Traditional manual counting (commonly targeting ~50 type I and ~75 type II fibers for adequate power^9^) is feasible for small datasets but is laborious and error-prone when biopsies yield hundreds to >1,000 fibers. Automated image analysis enables scalable, reproducible quantification of large cross-sections, yet existing tools^10–15^ often measure different endpoints and can be difficult to adapt across antibodies, imaging settings, or file formats.

Motivated by the above, in this paper, we developed a semi-automated image analysis pipeline for immunohistological images of skeletal muscle cross-sections that quantifies:

1. *Myonuclear number per muscle cell*,
2. *Pax7+ number per muscle cell*,
3. *Fiber type of each muscle cell*,
4. *Cross-sectional area of each muscle cell*,
5. *Perimeter of each muscle cell*.

This study is, to our knowledge, the first study to combine these five analytic targets into a single analysis pipeline compatible with standard widefield microscopy—a widely available platform in core facilities and individual laboratories. This workflow enables accurate, high-throughput quantification of muscle cross-sections while preserving full visual outputs for user review and quality control. The method has broad applicability in biomedical research and may help uncover mechanisms related to satellite cells and myonuclei within various physiological and pathological contexts across the lifespan.

## Methods and Results

### Ethical approval

The study was approved by the University of Oregon Institutional Review Board and conducted in accordance with *Declaration of Helsinki* ethical principles. All subjects gave informed written consent prior to study participation.

### Biopsies

Skeletal muscle biopsies were performed on human vastus lateralis in a distal to proximal sequence and in the fasted state. For serial biopsies the biopsy needle was oriented lateral from and perpendicular to the long axis of the femur with the biopsy needle window/opening situated in the mid-substance of the muscle tissue. Serial biopsies were thus obtained from the mid-substance of the vastus lateralis muscle that are 3 or more cm apart (and not angled)^13,14^. Biopsy samples were obtained using a Bergström needle with 5mm internal diameter with manual suction. Each biopsy yielded approximately 300mg of tissue (range ~100—500mg) with the fiber bundles approximating 2 to 7mm in length.

### Immunohistochemistry

Immediately following biopsy, tissue was trimmed of fat and connective tissue, aligned longitudinally, embedded in Optimal Cutting Temperature compound (OCT), and submerged into a section of liquid-phase isopentane (2-methylbutane) surrounded by frozen isopentane (freezing point: 160°C) achieved with liquid nitrogen and stored at −80°C. Tissue sections were cut at 7µm on a Leica CM 1850UV cryostat set to −25°C and air dried for one hour at room temperature protected from light.

Sections were fixed in ice cold acetone (stored at −20°C) for 3 minutes on the bench and washed in PBS. Sections were then blocked with 3% H_2_O_2_ for 7 minutes, washed in PBS, and blocked for 1 hour at room temperature in blocking buffer consisting of 2.5% to 5% normal goat serum (cat. no. 005-000-121, Jackson ImmunoResearch, West Grove, PA, USA), 2.5% to 5% normal donkey serum (cat. no. 017-000-121, Jackson ImmunoResearch, West Grove, PA, USA), and 1% glycine in PBS.

Primary antibodies for laminin (Figure 1A), type I myosin heavy chain (Figure 1C), and satellite cells (Figure 1D) were mixed in blocking buffer, allowed to incubate for 1 hour at room temperature, and then moved to 4°C for overnight incubation. Primary antibodies and concentrations were as follows: 1:75 BA-D5-c (deposited to the DSHB by Schiaffino, S., DSHB Hybridoma Product BA-D5), 1:200 Anti-Laminin (cat. no L9393, Sigma-Aldrich, Darmstadt, Germany), and 1:100 PAX7-c (deposited to the DSHB by Kawakami, A., DSHB Hybridoma Product PAX7).

**Figure 1.**
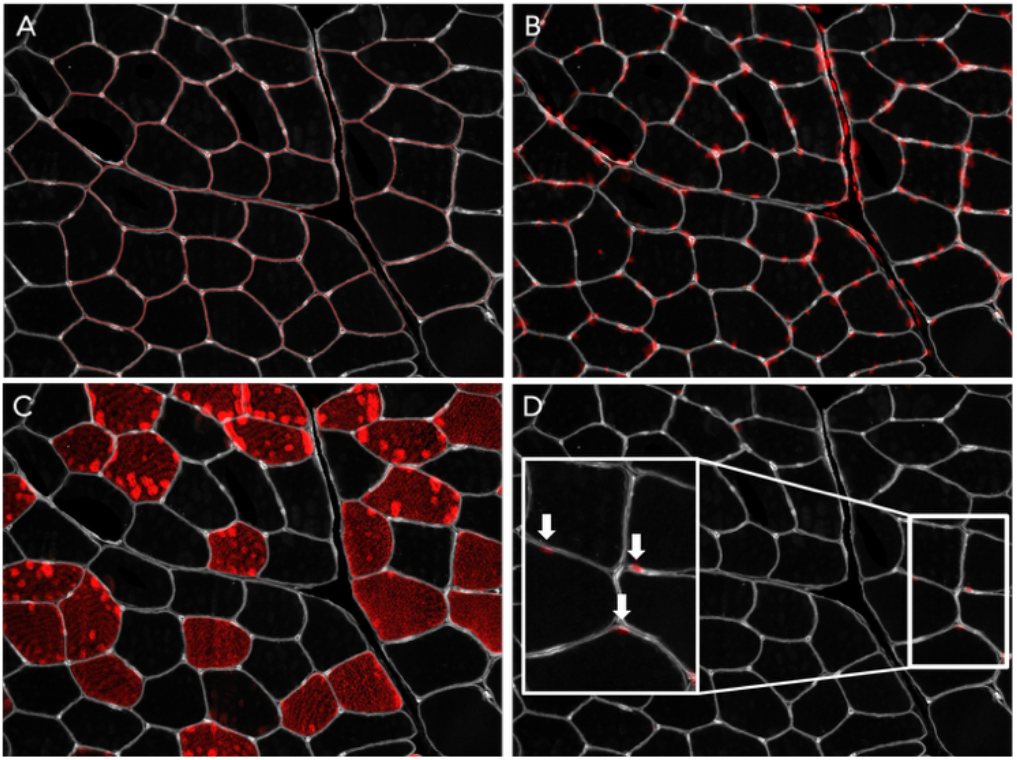
*Vastus Lateralis* cross-section histology. Laminin stain is shown in white, and parameters of interest are shown in red. Images taken at 20x. **(A)** Red overlay shows segmentation of myofibers by CellProfiler. **(B)** DAPI. **(C)** Myosin Heavy Chain I indicating type I fibers. **(D)** White arrows within callout indicate Pax7+ satellite cells.

After overnight incubation, sections were washed in PBS, blocked with 3% H_2_O_2_ for 7 minutes, and washed again in PBS. Secondary antibodies were mixed in blocking buffer and incubated for 1 hour at room temperature. Secondary antibodies and concentrations were as follows: 1:250 Cy3 IgG (cat. no. 715-165-150, Jackon ImmunoResearch, West Grove, PA, USA), 1:500 Alexa Fluor 647 IgG (cat. no. A21245, Invitrogen, Carlsbad, CA, USA), and 1:1000 Biotin-SP IgG (cat. no. 115-065-205, Jackon ImmunoResearch, West Grove, PA, USA). Following secondary incubation, sections were washed in PBS and incubated for 1 hour at room temperature in HRP-conjugated streptavidin (cat. no. B40932, Invitrogen, Carlsbad, CA, USA). Sections were then washed in PBS and incubated for 1 hour at room temperature in Alexa Fluor 488 (cat. no. B40932, Invitrogen, Carlsbad, CA, USA). Finally, sections were washed in PBS and mounted with SlowFade Diamond Antifade with DAPI (cat. no. S36964, Invitrogen, Carlsbad, CA, USA) (Figure 1B).

### Image acquisition

Tissue sections were imaged on a Leica DM4000B widefield fluorescence microscope with a Leica DFC360FX monochrome camera (pixel size 6.45 × 6.45 µm) and a Lumen200 metal halide lamp, (Prior Scientific, Cambridge, United Kingdom). Images were acquired at 20x magnification (HC PL FLUOTAR 20x/0.50; 1pixel = 1.9535µm) in four channels: Blue (Ex: 340-380nm, Em: 425nm), Green (Ex: 492nm, Em: 520nm), Yellow (Ex: 550nm, Em: 570nm), and Far-Red (Ex: 650nm, Em: 670nm). Exposure times, camera gain, and light intensity were adjusted for each channel to maximize dynamic range without saturation. These settings should be empirically determined by each user depending on imaging system. Multiple non-overlapping fields of view were collected systematically across each section.

### Image cleaning & pre-processing

To ensure the integrity of the automated analysis pipeline and maximize the reliability of downstream data, a standardized image “cleaning” protocol was implemented (Figure 2). The primary objective of this pre-processing stage was the systematic removal of optical artifacts and abnormal regions— such as overlapping or blurred laminin signals and out of focus or extraneous nuclei—that could potentially introduce noise or lead to false-positive identifications within the *CellProfiler* environment. The “cleaned” images better reflect the high-fidelity regions measured in manual, by-hand analysis.

**Figure 2.**
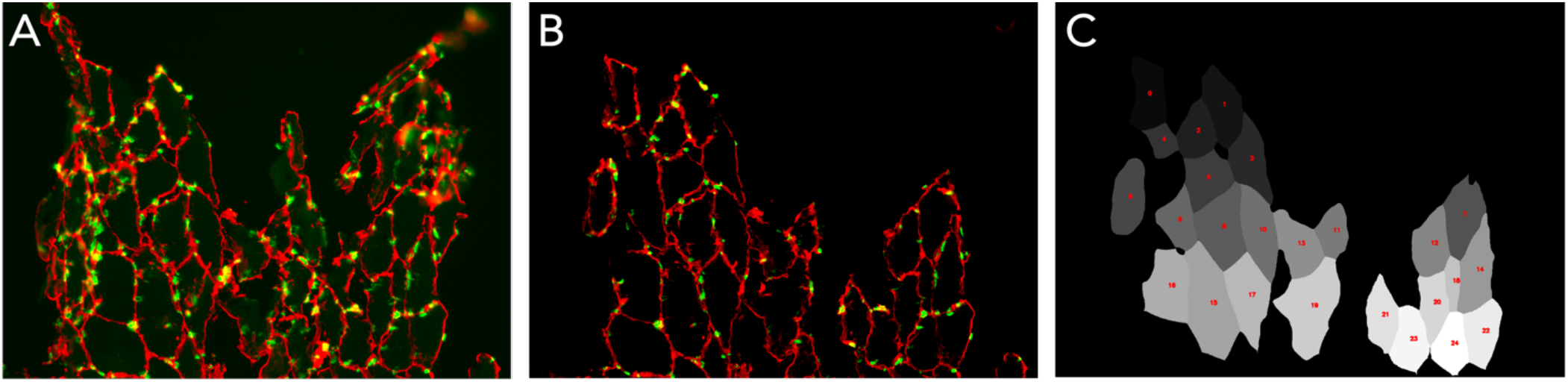
Imperfect images can be segmented by the program with pre-processing. Red staining shows laminin (cell borders) and green staining shows DAPI (nuclei). **(A)** Raw image overlay of the imperfect edge of a tissue section at 20x. **(B)** The same image overlay post-cleaning in FIJI. **(C)** Labeled cells identified by the pipeline from tissue section shown in A.

The workflow utilizes *FIJI/ImageJ* for manual curation and signal enhancement. Signal-to-noise ratio is first optimized through individual channel brightness and contrast adjustments; specifically, the “Minimum” and “Maximum” toggles were visually calibrated to enhance feature visibility.

The laminin, DAPI, and Pax7 channels are merged into a composite image using the “Merge Channels” function, ensuring “Keep source images” was enabled to preserve the underlying raw data. Areas identified as problematic— including non-specific staining, tissue folds, blood vessels, or poor nuclear definition—were manually circumscribed and removed using the “Clear” function.

Following artifact removal, the composite images were decomposed into their constituent channels (“Split Channels”) and exported as 8-bit .tif files. To ensure compatibility with the automated modular architecture of the *CellProfiler-Cellpose* integration, a strict naming convention was applied (e.g., Subject8_BL_L_Set1_Lam), facilitating the seamless batch processing and metadata assignment required for the final morphometric analysis.

### CellProfiler-Cellpose pipeline

We developed a custom *CellProfiler-Cellpose* pipeline to process cleaned immunohistochemistry images. This workflow automates 2D cellular and myonuclear segmentation, feature extraction, and file organization, yielding binarized mask outputs as well as visual overlays for muscle fibers, myonuclei, and satellite cells.

*CellProfiler* (v4.2.8) is an open-source software tool for automated image analysis, currently developed and maintained in Cimini Lab at the Broad Institute of MIT and Harvard^18^. *CellProfiler* is designed for high-throughput, quantitative microscopy workflows, using modular pipelines for object segmentation and measurement.

The pipeline integrates *Cellpose* (v2.3.2), a machine learning model plugin developed by Carsen Stringer and Marius Pachitariu at the Howard Hughes Medical Institute (HHMI) Janelia Research Campus. *Cellpose* uses a U-Net–style convolutional neural network to predict likely cell regions and to reconstruct cell boundaries.

*Cellpose* was selected for myonuclear segmentation as its pretrained models accurately segment diverse tissue morphologies without the need for custom training. This compatibility facilitates the automated segmentation of irregular and overlapping objects within a high-throughput framework. Further information on the *Cellpose* algorithm can be found here: https://doi.org/10.1038/s41592-020-01018-x.

The pipeline operates across two computing environments: a standard Windows desktop workstation and a high-performance supercomputer. Deployment occurs either through Docker (v28.2.2) or within a local Python environment. Docker provides enhanced stability on desktop systems, while native Python execution yields superior performance on high-performance clusters. For further computing specifications, see supplemental.

Careful reading of attached supplemental documents and the *CellProfiler* manual (https://cellprofiler-manual.s3.amazonaws.com/CellProfiler-4.2.8/index.html) is encouraged before using the pipeline. As highlighted in the manual, for each module within the CellProfiler pipeline there are several adjustable variables that will impact the way the module functions. Below, we have listed the values that were found to be most accurate for our images within this pipeline, but users should test and adjust these variables to tailor it to their images specifically.

#### Overview of the CellProfiler pipeline

##### 1. Metadata extraction & image organization

The *Metadata* module utilizes Regular Expression patterns to dynamically group and extract metadata, including Subject ID, biopsy timepoint, leg (L/R), image set, and staining protocol. The *NamesAndTypes* module identifies specific stains using the following tags:

- *Laminin*: laminin
- *DAPI*: nuclei
- *Pax7*: satellite
- *MyHC1*: myhc1

The *Groups* module then organizes input image sets by *“Subject”, “Day”, “Side”*, and *“Set”*, as extracted by the *Metadata* module.

##### 2. Cellular and myonuclear segmentation

For cellular segmentation, the pipeline employs the ‘cyto2’ detection mode within the *Cellpose* module on laminin-stained images to identify sarcolemmal boundaries (Figure 3). For myonuclear segmentation, the workflow initiates with a *GaussianFilter* (sigma = 3) to smooth the DAPI signal and reduce noise. To address the high sensitivity of the default *Cellpose* ‘nuclei’ model, the pipeline implements an iterative subtraction logic to minimize false-positive detections:

1. *Initial Detection*: A *RunCellpose* module invokes the ‘nuclei’ model to identify primary nuclear candidates (flow_threshold = 0.4; cellprob_threshold = 0.0).
2. *Background Normalization*: The *MeasureObjectIntensity* module calculates the current object intensity distribution, and the *ImageMath* module subtracts the 17th percentile of pixel intensities from the original DAPI signal. This suppresses low-level background fluorescence and non-specific staining.
3. *Refined Segmentation*: The cleaned image passes through a second *RunCellpose* module with optimized parameters to ensure precise boundary adherence (Figure 4).

**Figure 3.**
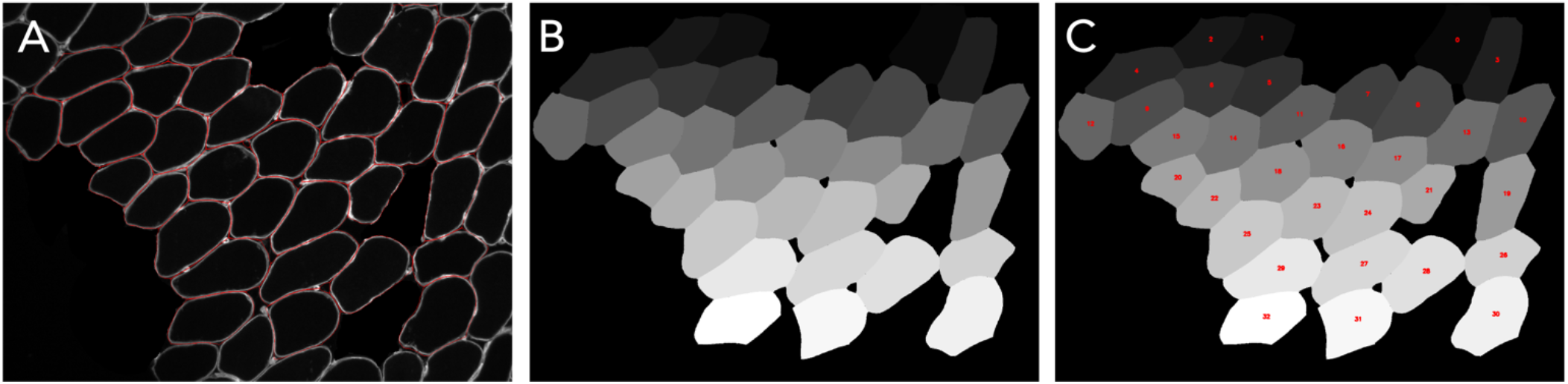
Myofiber segmentation with *Cellpose* ‘cyto2’ detection model. **(A)** White staining shows fluorescent laminin signal, with red overlay showing cells accepted by the pipeline. This overlay is saved for each image set to allow for manual quality control. **(B)** Grayscale object masks generated and saved by the pipeline for use in the Python script. **(C)** Masks identified by the pipeline are numbered by the Python script so that each cell within each image has a unique identifier, allowing for manual quality controls.

**Figure 4.**
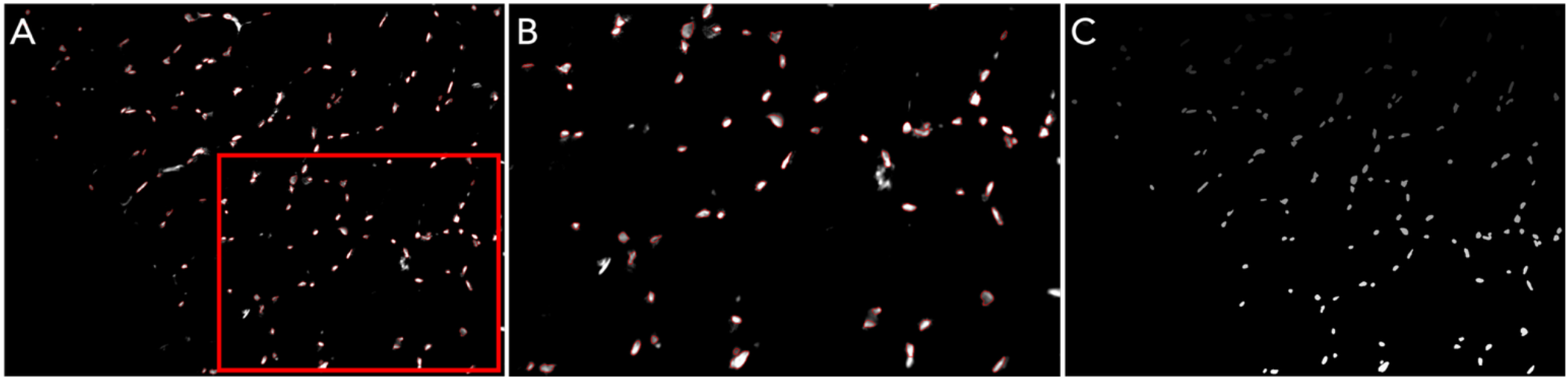
Nuclear segmentation with *Cellpose* ‘nuclei’ detection model. **(A)** White staining shows fluorescent DAPI signal, with red overlay showing nuclei accepted by the pipeline. This overlay is saved for each image set to allow for manual quality control. **(B)** Callout from red box in (A). **(C)** Grayscale object masks generated and saved by the pipeline for colocalization with myofibers in the Python script.

##### 3. Satellite cell segmentation

Pax7+ satellite cell segmentation utilizes the *IdentifyPrimaryObjects* module, as deep-learning architectures prove insufficient for resolving sparse Pax7 signals against the sarcoplasmic background (Figure 5). The module employs an Otsu thresholding strategy to isolate satellite cells within a defined pixel diameter range of 16-60; this parameter can be adjusted based on the expected size of objects of interest. Objects are further filtered by circularity and intensity to exclude signal artifacts.

**Figure 5.**
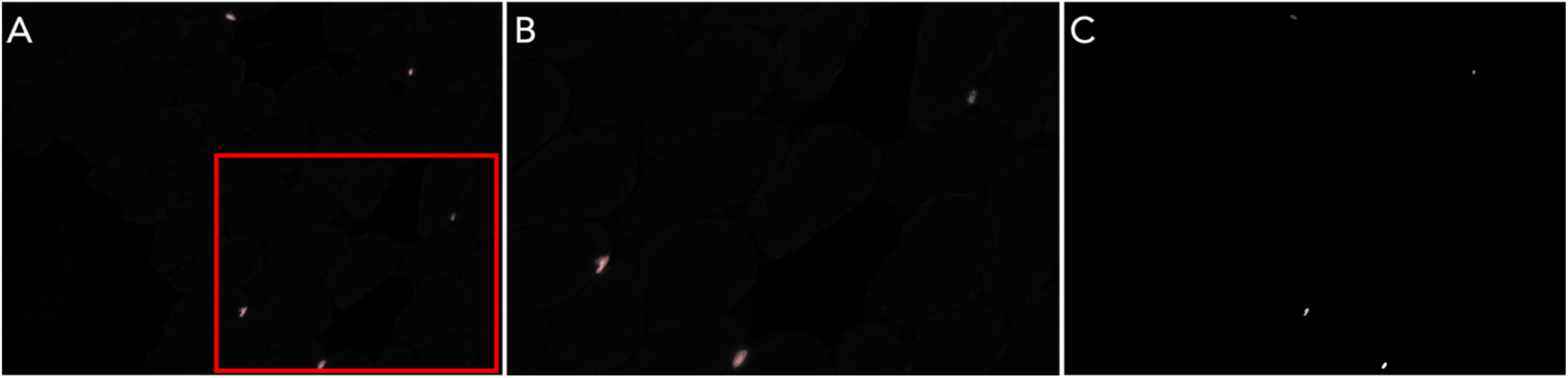
Satellite Cell segmentation with *IdentifyPrimaryObjects* model. **(A)** White staining shows fluorescent Pax7+ signal, with red overlay showing satellite cells accepted by the pipeline. This overlay is saved for each image set to allow for manual quality control. **(B)** Callout from red box in (A). **(C)** Grayscale object masks generated and saved by the pipeline for colocalization with myofibers in the Python script.

The pipeline is configured for high-throughput batch processing. By leveraging the *Metadata* and *Groups* modules, the system systematically organizes segmentation masks and morphological data into a standardized hierarchical directory structure. This consistent naming convention allows a downstream Python-based analysis script to programmatically parse and aggregate data for final statistical refinement.

### Python-based processing

To transition from raw segmentation masks to refined, biologically relevant datasets, we developed a custom Python-based processing pipeline utilizing the OpenCV, pandas, and NumPy libraries. The pipeline is designed to automate the integration of multi-channel spatial data, specifically matching nuclear and satellite cell positions to individual muscle fiber domains. This stage of the workflow (1) ensures objective fiber-type assignment, (2) quantifies morphological parameters in physical units, and (3) aggregates data at the biopsy level while maintaining full traceability of individual fibers.

#### 1. Input Structure and Data Parsing

The pipeline employs a dynamic directory parser that identifies image sets based on a standardized naming convention. The script scans designated subdirectories for four critical components:

- *Laminin Masks*: Grayscale images where each fiber is represented by a unique pixel intensity value.
- *Nuclei Masks*: Binary/grayscale masks representing DAPI-stained nuclei.
- *Satellite Cell Masks*: Binary/grayscale masks representing Pax7-stained cells.
- *MyHC1 Images*: Raw fluorescence images used for fiber-type intensity quantification.

#### 2. Grayscale Mask Decomposition

Because *Cellpose*’s segmentation outputs a single label matrix where each object has a unique grayscale value, the function *split_mask_by_gray_level* was implemented. This function iterates through the unique pixel values of the image, extracting each identified fiber or nucleus into an individual binary mask. This decomposition is essential for the subsequent bitwise overlap analysis and morphological measurements.

#### 3. Fiber-Object Assignment

To accurately count nuclei and satellite cells per fiber, we developed a “greatest overlap” assignment logic via the *overlap_with_objects* and *fiber_assign* functions.

- *Bitwise Intersection*: For every fiber mask, the pipeline performs a bitwise AND operation against all identified object masks (nuclei or satellite cells).
- *Primary Assignment*: To account for objects that may partially overlap with multiple fibers, each object is strictly assigned to the fiber with which it shares the largest number of overlapping pixels.
- *Quantification*: This results in a discrete count of myonuclei and satellite cells localized specifically to each fiber’s cytoplasmic domain.

#### 4. Morphometry and Scaling

Fiber area and perimeter are calculated using the *calculate_area_and_perimeter* function. To ensure data is reported in biological units, a pixel-to-micron scaling factor is applied (1.9535 pixels/micron for 20x magnification).

#### 5. Fiber Type Assignment

The pipeline automates the classification of type I and type II fibers using MyHC1 fluorescence. Background is removed from images with the function *background_subtract* using the classic rolling ball algorithm (ball radius = 105 px; chosen empirically to exceed the largest MyHC1+ fiber), and this radius should be adjusted to match cell size in other datasets. The function *calculate_average_intensity* computes the mean pixel intensity from the background-subtracted image specifically within the coordinates defined by each fiber mask. Fibers with a mean MyHC1+ intensity >20 are classified as type I (assigned value: 1) and those below are type II (assigned value: 0). The threshold value was derived from an intensity analysis of 3,722 cells from 12 human biopsies within our data set (Figure 6).

**Figure 6.**
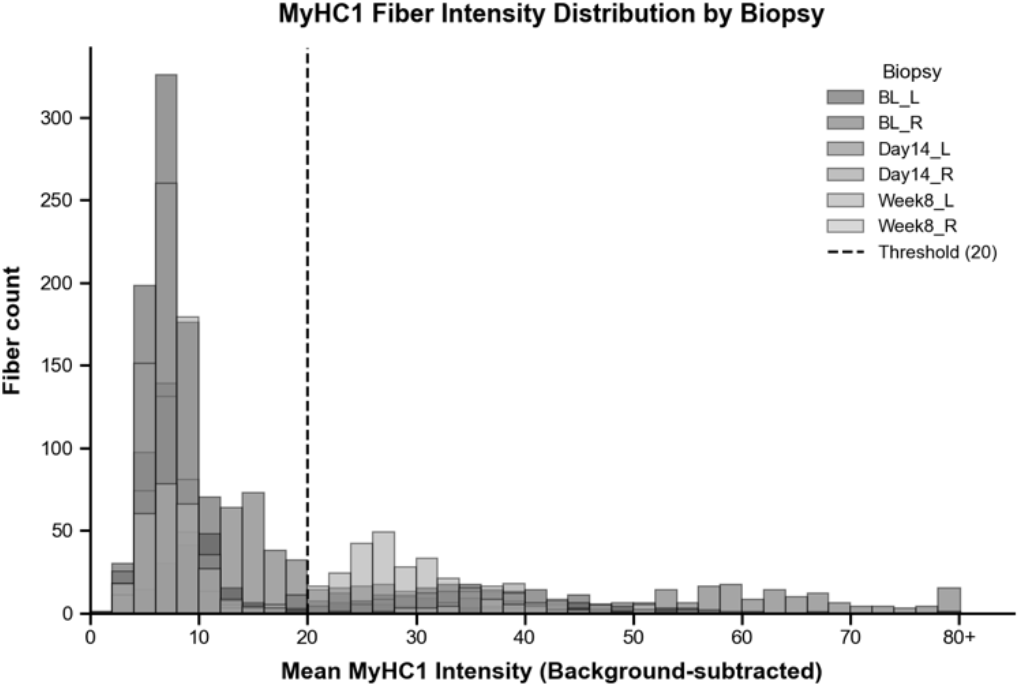
Intensity threshold for automated fiber typing. Analysis of intensities within segmented myofibers from 3,722 muscle cells from 12 human biopsies was used to establish a threshold between type I and type II fibers. The frequency distribution shows two groups, with type II fibers being the group below an intensity of 20 and type I fibers being the group above an intensity of 20. The dotted line represents the threshold at pixel value 20.

#### 6. Batch Orchestration & Outputs

The *process_subject* function automates the workflow across all timepoints and legs for a given subject. The pipeline generates several high-value outputs:

- *Excel Workbooks*: A unified .xlsx file containing sheets for each biopsy, including columns for fiber ID, area, perimeter, nuclei count, satellite cell count, and MyHC1 intensity.
  ∘ Certain identified cells, specifically those labeled as cell number “0”, should be sorted and removed from the data set as the pipeline assigns extraneous nuclei to these cells. Any cells that had 10 or more nuclei assigned to it are manually verified or removed from the spreadsheet.
- *Diagnostic Overlays*: Labeled laminin images are generated for rapid visual quality control (QC) to verify the accuracy of the automated counts.

### Validation of pipeline outputs

For pipeline validation, 310 cells were matched and analyzed both manually and by the pipeline. For each cell, borders were traced by hand and myonuclei, Pax7+ satellite cells, and fiber type were scored manually^19^. Area and perimeter of each cell were recorded based on hand drawn tracings. The pipeline produced the same five metrics for the 310 cells. Methods were compared using Bland Altman analyses measuring the absolute difference, paired two-tailed t-tests (α = 0.05), frequency distributions, and correlation analyses.

Bland Altman comparison of manual versus pipeline myonuclei per cell resulted in a bias of −0.003 nuclei with 95% limits of agreement of −2.831 to 2.824 nuclei (Figure 7A). Across 310 cells, the average manual myonuclei/cell was 2.471 and the average pipeline myonuclei/cell was 2.474, with no significant difference by paired t-test (p=0.9686; Figure 7B). Frequency distributions for both methods are shown in Figure 7C. The two methods were moderately correlated (r=0.4926, p<0.0001).

**Figure 7.**
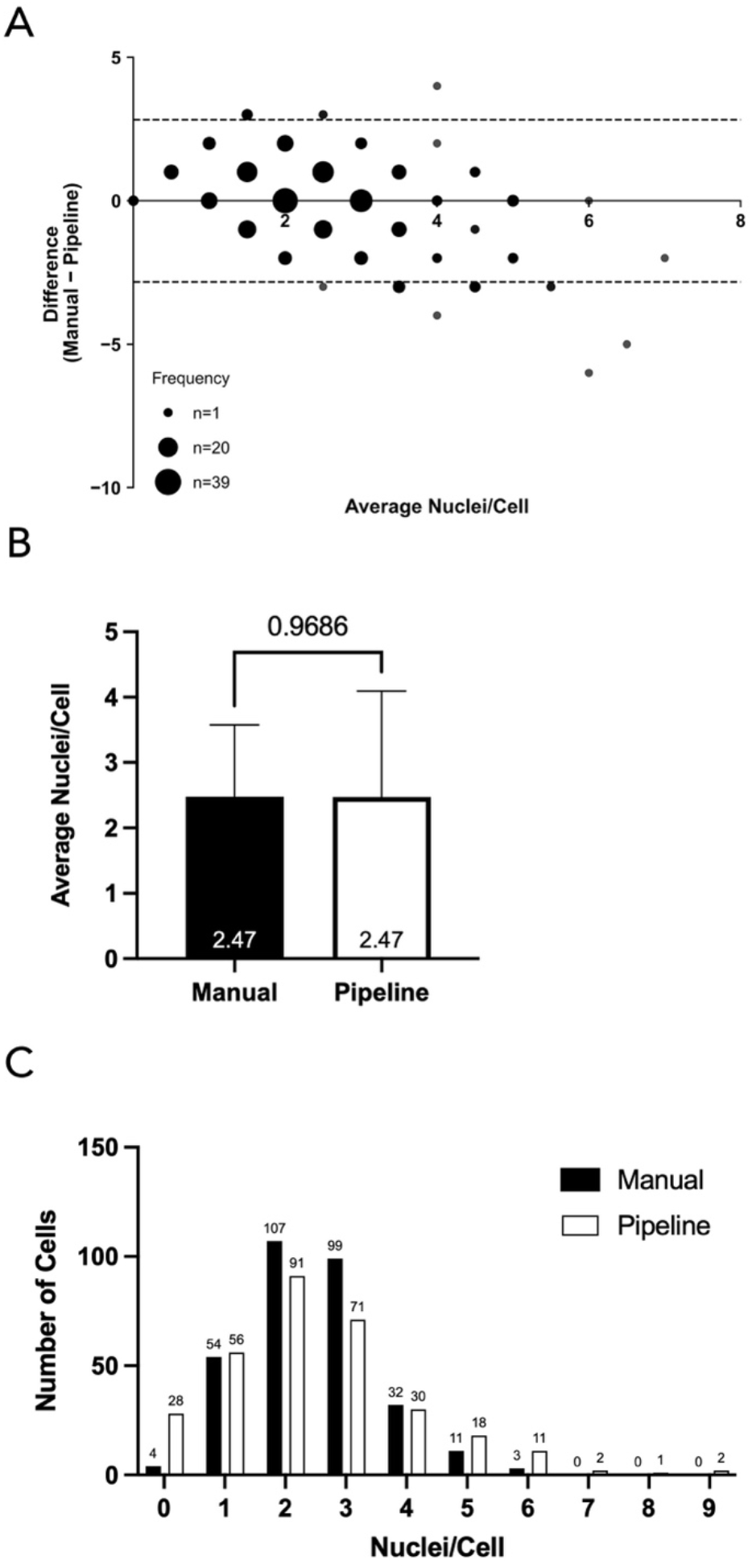
Comparison of nuclei/cell counted manually versus by the pipeline (N=310) **(A)** Bland Altman 95% limits of agreements plot comparing manual nuclear counting to pipeline nuclear counting. The frequency of data points is represented by point size. **(B)** Average nuclei/cell as counted by manual analysis and by the pipeline. Paired t-test resulted in a two-tailed p-value of 0.9686. Data is presented as mean ± standard deviation. **(C)** Frequency distribution of nuclei/cell for manual (black bars) and pipeline (white bars) analysis.

Bland Altman comparison of manual versus pipeline for Pax7+ satellite cells per cell resulted in a bias of −0.0097 satellite cells with 95% limits of agreement of −0.5441 to 0.5247 satellite cells (Figure 8A). Across the 310 cells, the manual average satellite cells/cell was 0.110 and the pipeline average satellite cells/cell was 0.119, and a paired t-test between the two methods revealed a p-value of 0.5325 (Figure 8B). The frequency distribution shows the range of satellite cells/cell for both the manual and the pipeline methods (Figure 8C). The two methods were moderately correlated (r=0.6762, p<0.0001).

**Figure 8.**
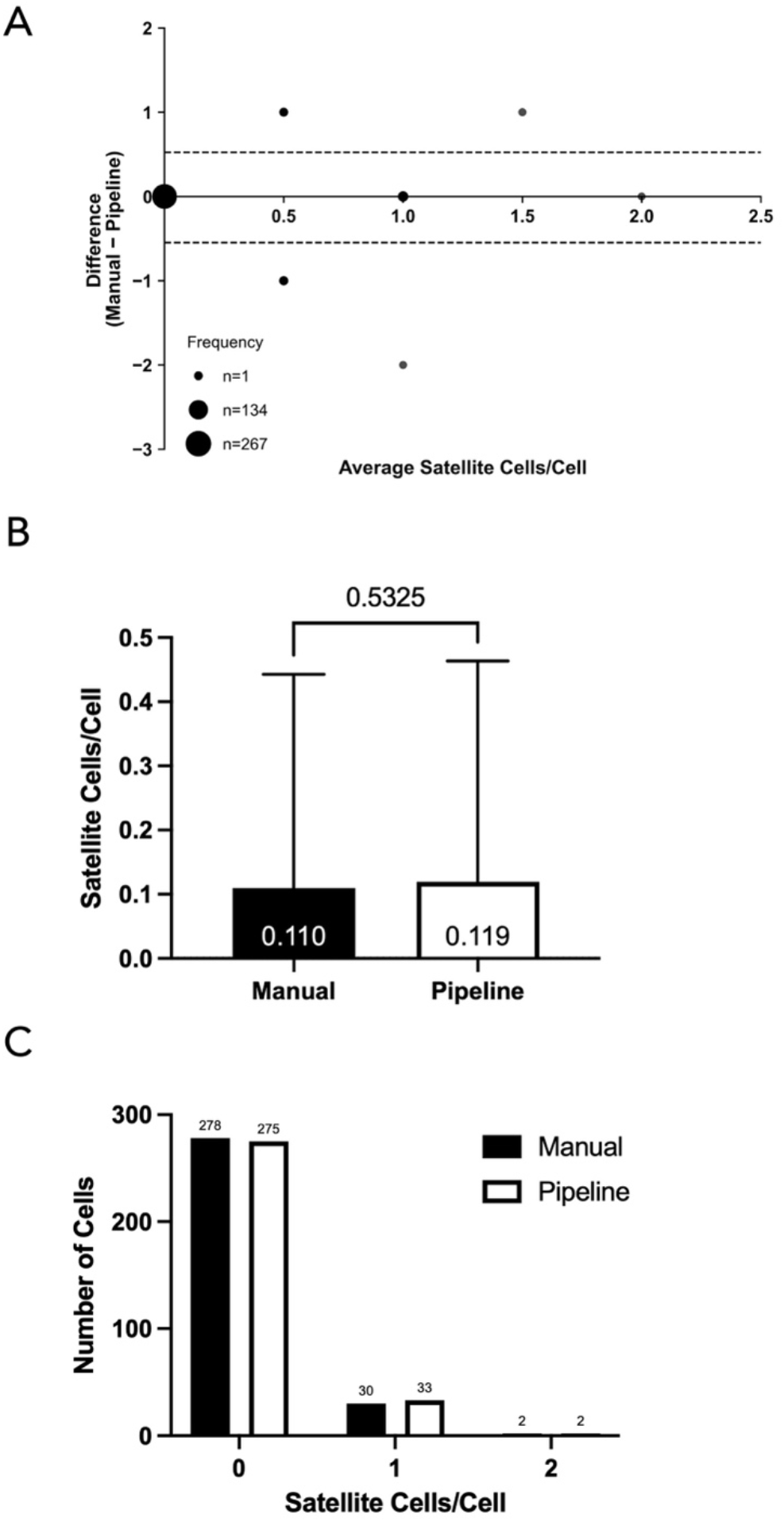
Comparison of satellite cells/cell counted manually versus by the pipeline (N=310) **(A)** Bland Altman 95% limits of agreements plot comparing manual satellite cell counting to pipeline satellite cell counting. The frequency of data points is represented by point size. **(B)** Average satellite cells/cell as counted by manual analysis and by the pipeline. Paired t-test resulted in a two-tailed p-value of 0.5325. Data is presented as mean ± standard deviation. **(C)** Frequency distribution of satellite cells/cell for manual (black bars) and pipeline (white bars) analysis.

For fiber typing, Bland Altman analysis resulted in a bias of −0.0032 type I cells with 95% limits of agreement of −0.2525 to 0.2460 (Figure 9A). Across 310 cells, the manual proportion of type I fibers was 0.155 manually versus 0.158 for the pipeline, and the paired t-test revealed a p value of 0.6555 (Figure 9B). Manual fiber typing identified 48 type I fibers and 262 type II fibers while automated fiber typing identified 49 type I fibers and 261 type II fibers (Figure 9C). The two methods were highly correlated (r=0.9390, p<0.0001).

**Figure 9.**
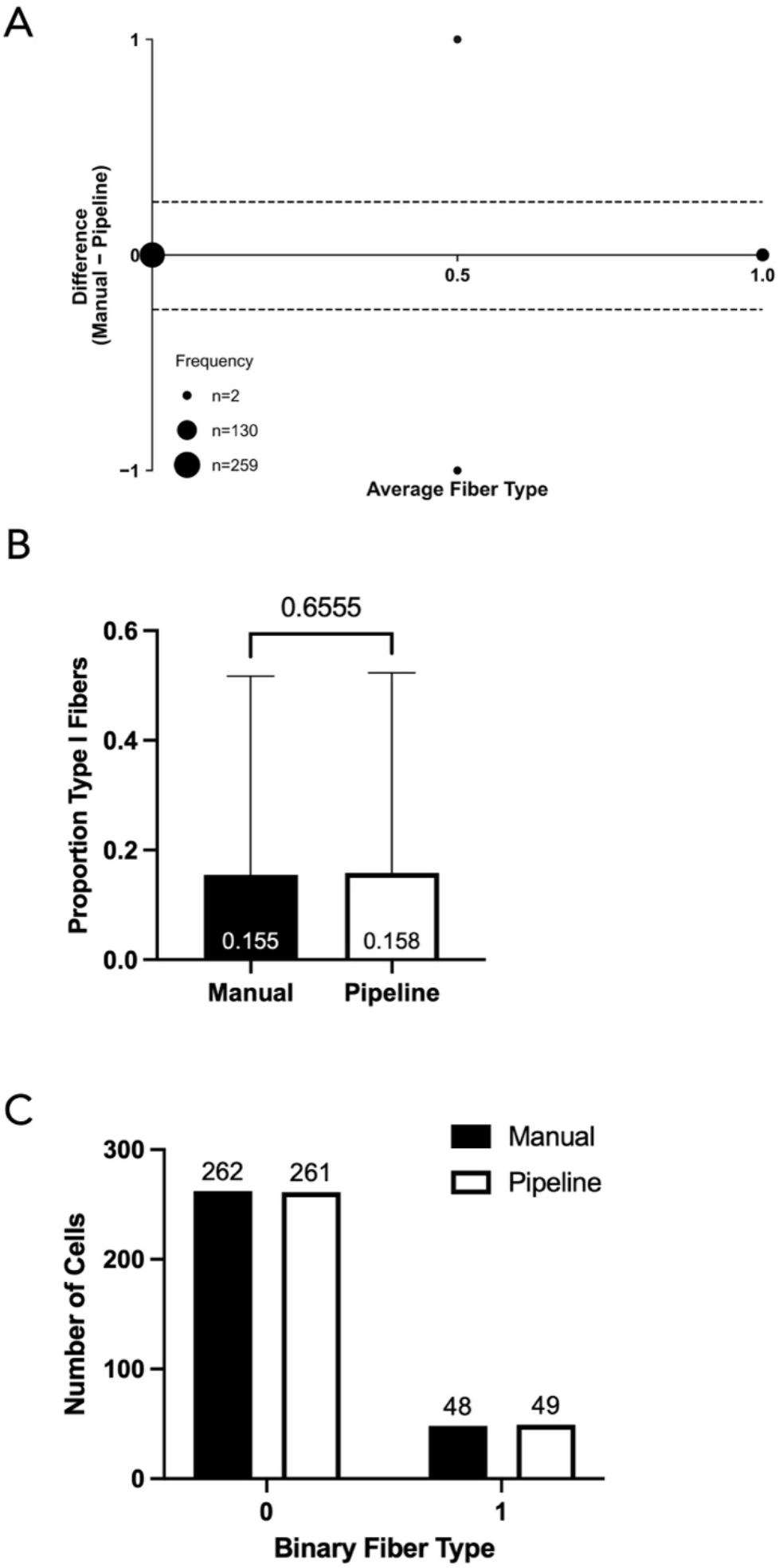
Comparison of fiber type assessed manually versus assessed by the pipeline (N=310) **(A)** Bland Altman 95% limits of agreements plot comparing manual fiber typing to pipeline fiber typing. The frequency of data points is represented by point size. **(B)** Proportion of type I fibers as counted by manual analysis and by the pipeline. Paired t-test resulted in a two-tailed p-value of 0.6555. Data is presented as mean ± standard deviation. **(C)** Frequency distribution of type I and type II fibers for manual (black bars) and pipeline.

To validate area measurements, the manual outlines measured by FIJI were compared to automated area outputs from CellProfiler. Bland Altman analysis comparing manual and pipeline cross-sectional area (CSA) measurements showed a bias of −527µm^2^ with 95% limits of agreement of −1072 to 18.11µm^2^ (Figure 10A). Across the 310 cells, the mean manual CSA was 3005µm^2^ versus 3532µm^2^ for the pipeline, and the paired t-test was significant (p-value < 0.0001; Figure 10B). Frequency distributions of CSA for both methods are shown in Figure 10C. The methods were highly correlated (r=0.9592, p<0.0001; Figure 10D).

**Figure 10.**
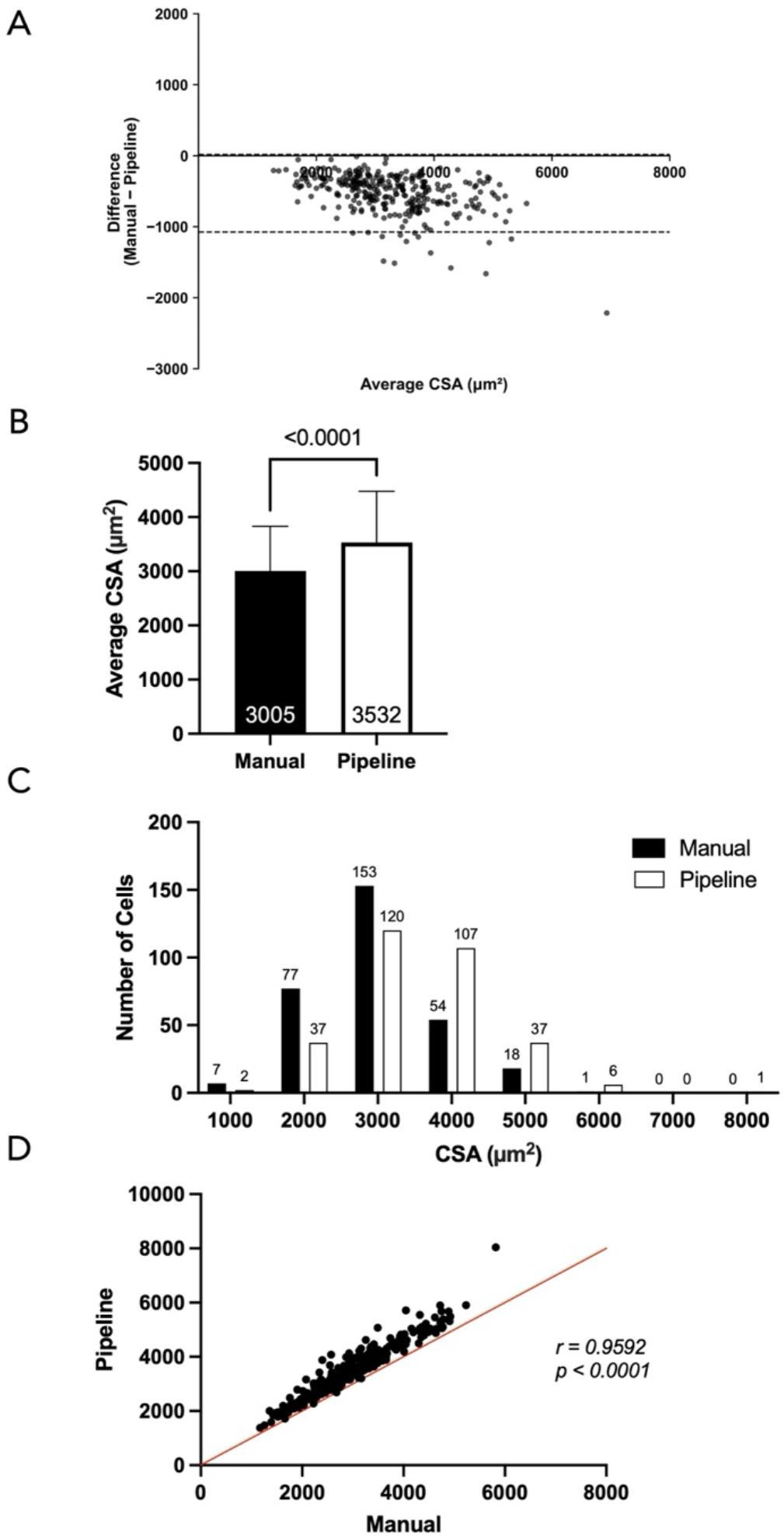
Comparison of cross-sectional area (CSA) measured manually versus by the pipeline (N=310) **(A)** Bland Altman 95% limits of agreements plot comparing manual cross-sectional area to pipeline cross sectional area. **(B)** Average cross-sectional area counted by manual analysis and by the pipeline. Paired t-test resulted in a two-tailed p-value less than 0.0001. Data is presented as mean ± standard deviation. **(C)** Frequency distribution of cross-sectional area for manual (black bars) and pipeline (white bars) analysis. **(D)** Correlation between cross-sectional area from the pipeline and measured manually; r = 0.9592 and p < 0.0001.

For perimeter validation, the perimeters of the manually drawn outlines used for CSA were measured and recorded by Python and compared to automated perimeter outputs from CellProfiler. Bland Altman comparing manual versus pipeline for perimeter resulted in a bias of −15.13µm with 95% limits of agreement of −41.69µm to 11.43 µm (Figure 11A). Across the 310 cells, the manual average perimeter was 249µm versus 264µm for pipeline, and the paired t-test was significant (p<0.0001; Figure 11B). Frequency distributions are shown in Figure 11C. The methods were highly correlated (r=0.9354, p<0.0001; Figure 11D).

**Figure 11.**
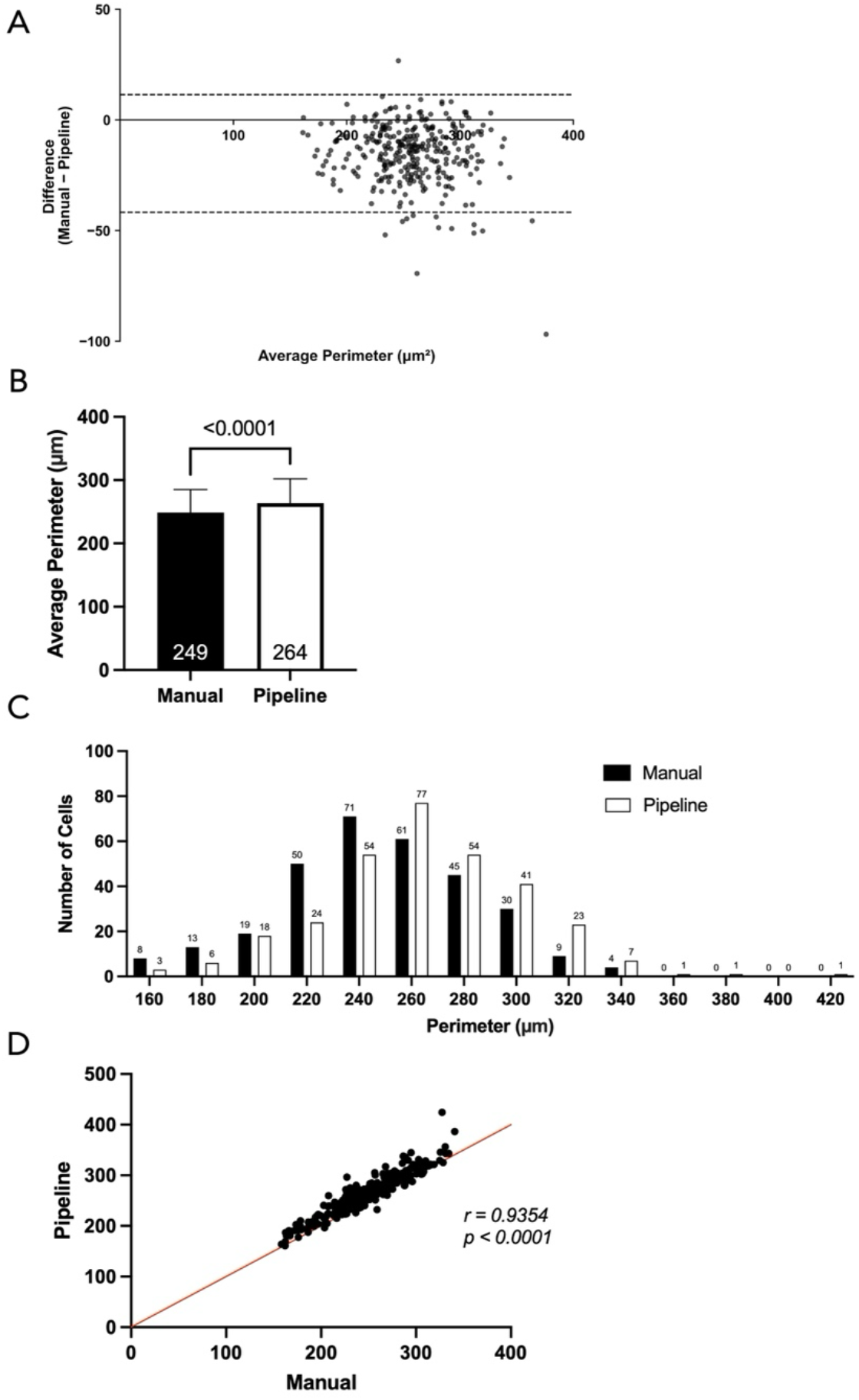
Comparison of perimeter measured manually versus by the pipeline (N=310) **(A)** Bland Altman 95% limits of agreements plot comparing manual perimeter pipeline perimeter. **(B)** Average perimeter counted by manual analysis and by the pipeline. Paired t-test resulted in a two-tailed p-value less than 0.0001. Data is presented as mean ± standard deviation. **(C)** Frequency distribution of perimeter for manual (black bars) and pipeline (white bars) analysis. **(D)** Correlation between perimeter from the pipeline and measured manually; r = 0.9354 and p < 0.0001.

## Discussion

In this study, we developed and validated a semi-automated pipeline analysis method to quantify Pax7+ satellite cells, myonuclei, cross-sectional area, and perimeter by fiber type in skeletal muscle cross-sections. This approach delivers reproducible, unbiased measurements with considerably reduced analysis time and user-dependent compared with manual analysis.

While several automated pipelines exist for analyzing muscle tissue ultrastructure^10–15^, none contain our parameters of interest— Pax7+ satellite cells, myonuclei, fiber CSA, perimeter, and fiber type—into a single workflow. Additionally, existing pipelines are dependent on images that are free of extraneous (background) signals or imperfect tissue regions^10^. Our goal was to build a robust, modular pipeline that quantifies these specific endpoints, while remaining modular enough to be adapted to other tissue types or parameters of interest. For example, within our lab, the pipeline presented here was readily modified to identify macrophages (in place of satellite cells) and atrophied myofibers from an elderly clinical cohort.

Our first aim was accurate quantification of myonuclei per myofiber, which can be difficult due to other cell types present within the muscle cross-section. By combining FIJI-based pre-processing to remove extraneous signal with automated Python analysis, the pipeline yields reproducible nuclear counts that closely match manual scoring. Although the FIJI image cleaning step in is the most time-consuming component, it is essential for reducing false positives and improving confidence in downstream outputs. Overall, the pipeline workflow is substantially faster and far more reproducible than manual analysis.

Our second aim was to achieve accurate quantification of Pax7+ satellite cells per fiber. This was achieved using the highly modular CellProfiler platform. By specifying input size, circularity, and intensity filters tuned to Pax7+ signal, the pipeline reliably discriminated satellite cells from background artifacts.

The third aim was to scale-up throughput to quantify hundreds to thousands of muscle cells per cross-section. Published power estimates reveal a lower limit of 50 type I and 75 type II fibers are required for comparisons of satellite cells per myofiber^9^, we selected a minimum threshold of 150 cells per biopsy. While dependent on biopsy size, our pipeline can process over 1000 cells per sample, substantially increasing sample representation and efficiency.

The pipeline offers several practical advantages beyond its main outputs. CellProfiler’s modular design allows for the easy addition of other parameters or the adjustment of current parameters to accommodate different tissue types, antibodies, or imaging conditions^11^. The pipeline generates and saves segmentation overlays for each parameter within each image set, and the custom Python script assigns unique identifiers to every cell, features that are crucial for quality control and auditing. Finally, the Python script compiles results into a single Excel file that can be easily sorted for statistical analysis.

## Limitations

While the primary outputs (myonuclei/cell, Pax7+ satellite cells/cell, and fiber type) closely match manual measurement, systematic differences remain for CSA and perimeter. Manual outlines were drawn directly on or slightly within the laminin border, using a 10-pixel brush in FIJI (Figure 12A and 12C). In contrast, the pipeline places segmentation on or slightly outside of the laminin border, with a 1-pixel outline (Figure 12B and 12D). Because the CSA and perimeter are computed from inside the drawn outline, these small boundary offsets produce measurable differences in calculated area and perimeter. Similar discrepancies between manual and automated fiber outlines have been previously reported^10^.

**Figure 12.**
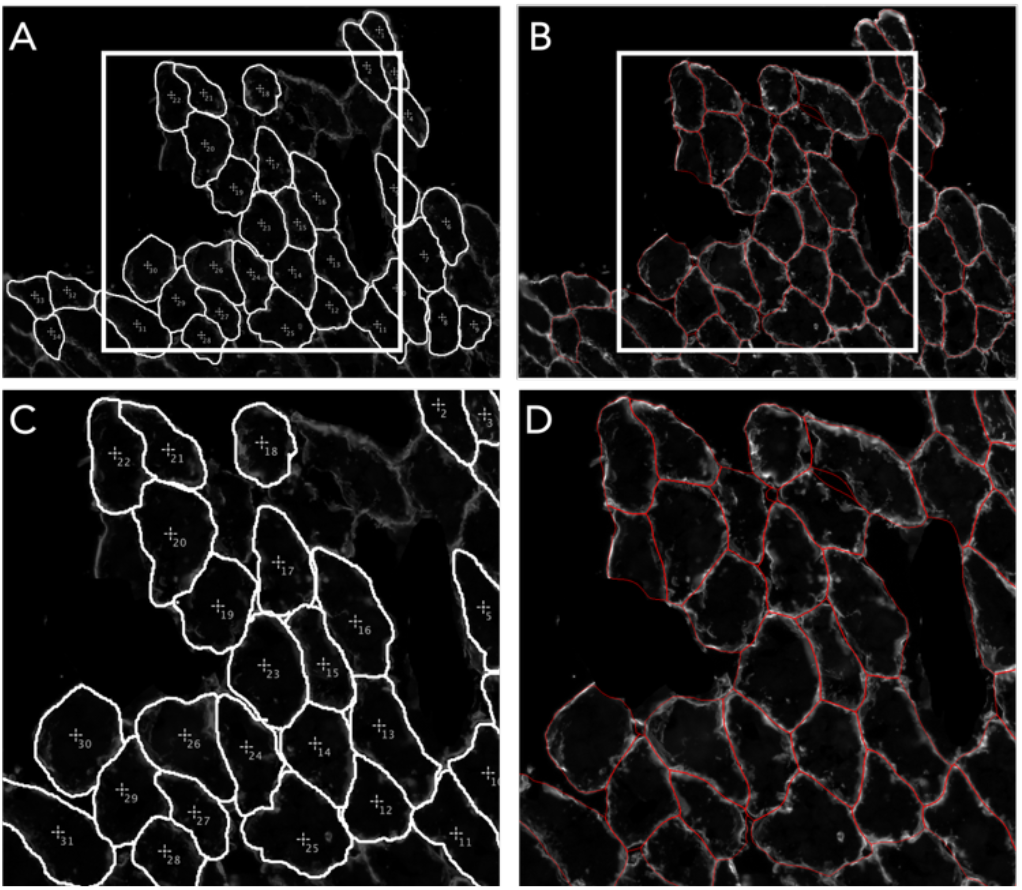
Manual versus CellProfiler cell outlines. **(A)** Manually drawn outlines shown in white, drawn over the raw laminin stain. Brush width used in FIJI was 10 pixels. **(B)** Automated outlines shown in red, drawn over the raw laminin stain shown in (A). Line width used by CellProfiler was 1 pixel. **(C)** Callout from white box in (A). **(D)** Callout from white box in (B).

Other limitations are the pipeline requires image cleaning and strict file naming before analysis. Although time consuming, removing extraneous artifacts and ensuring clear target visibility substantially improves the accuracy of the final output. Similarly, the strict file naming allows for automated organization of and batch processing of large image sets. As noted above, fibers labeled “fiber 0” include extraneous nuclei not assigned to any fiber—and these cases should be visually verified or excluded prior to analysis.

Finally, the *RunCellpose* “nuclei” model may fail to detect objects in images with very sparse or very dim DAPI signal, returning zero nuclei for those fields; this is generally when <10 cells occupy the field of view. The issue can also occur if the DAPI signal is very dim. This issue has been reported in community forums (e.g., GitHub, Image.sc). In our experience sparse images are uncommon, but, if dim, sparse, or noisy images occur frequently, users can try alternative detection modes within *RunCellpose*, or upgrade to the newer version of Cellpose, (e.g., Cellpose3) to improve detection.

In conclusion, we present a semi-automated pipeline that, (1) quantifies myonuclear number per fiber; (2) quantifies Pax7+ satellite cells per fiber; (3) quantifies CSA and perimeter per fiber; (4) assigns fiber type, and (5) aggregates outputs for quality control and ease of statistical analysis. Designed and optimized for batch processing and organization of large numbers of widefield image, the workflow generates rich datasets describing myonuclear and satellite cell dynamics across entire cross-sections. This workflow has the potential for broad application in muscle biology and clinical settings alike.

## Acknowledgements

We would like to acknowledge the Genomics & Cell Characterization Core Facility at the University of Oregon. We thank members of the Dreyer Lab for comments and suggestions while testing our pipeline and drafting this manuscript.

## Graphical abstract

*Created in BioRender. Megowan, H. (2026)* https://BioRender.com/9457v5e

## Author contributions

Conceptualization: H.G.M., M.L., A.S., H.C.D. Methodology: H.G.M., M.L., A.S. Investigation: H.G.M., M.L., K.B.A. Visualization: H.G.M., M.L., K.B.A. Funding Acquisition: H.C.D. Project Administration: H.G.M., H.C.D. Supervision: A.C.F., J.S., H.C.D. Resources: A.C.F., H.C.D. Writing – original draft: H.G.M., M.L. Writing – review & editing: H.G.M., A.C.F., J.S., H.C.D.

## Data and materials availability

All data and materials are available in the manuscript and supplemental protocol. Code is available at: https://github.com/DreyerLabUO/23Wu-CrossSection.

## Supplemental Documentation

## I. Image Acquisition & Pre-Processing

### A) Image processing with FIJI/ImageJ

#### Setup

1. Install “*FIJI with Batteries”* from web browser.

#### Procedure

##### 1. Image Acquisition and Initial Standardization

The dataset for each sample must consist of four channels: Laminin, DAPI, MyHC1, and Pax7.

1. Selection: Evaluate all available images. If duplicate DAPI channels exist, select the image with the highest signal-to-noise ratio. If neither is superior, both may be retained for later merging.
2. File Formatting: All images must be saved in .tif format.
3. Preliminary Renaming: Before processing in FIJI, rename files to ensure compatibility with the *CellProfiler* pipeline. Alternative nomenclatures can be used, but will require modification of metadata parsing pattern in *CellProfiler* pipeline.
  ∘ Remove technical strings (e.g., replace _ch00 with a blank space).
  ∘ Abbreviate “Baseline” to BL and “Laminin” to Lam.
  ∘ Standard Nomenclature: SubjectID_Timepoint_Side_Set#_Channel (e.g., Subject8_BL_L_Set1_Lam).

##### 2. Preparation of the MyHC1 Channel

The MyHC1 image serves as a reference and does not require manual cleaning.

- Rename the MyHC1 file according to the protocol in Step 1.
- Transfer the file directly to the designated “Cleaned Images” output directory.

##### 3. Contrast Adjustment and Channel Merging (FIJI/ImageJ)

Import the Laminin, Pax7, and DAPI images into FIJI.

1. Optimization: If necessary, adjust visualization parameters via Image > Adjust > Brightness/Contrast. Modify the minimum and maximum sliders to enhance feature definition. These adjustments are for visualization during cleaning and do not require separate saving.
2. Bit-Depth Verification: Ensure all channels are consistent (e.g., 8-bit). If a “bit-depth” error occurs, convert the image via Image > Type > 8-bit.
3. Channel Merging: To facilitate simultaneous cleaning across channels, merge the images via Image > Color > Merge Channels.
  ∘ Assign arbitrary colors to each channel based on visual clarity.
  ∘ Crucial: Enable the “Keep source images” option to preserve the original single-channel data.

##### 4. Manual Artifact Removal (“Cleaning”)

Artifacts, such as tissue folds or non-specific staining, must be removed to prevent errors in downstream segmentation.

1. Selection: Use the Freehand Selection Tool to circumscribe the artifact.
2. Deletion: Navigate to Edit > Clear to remove the signal within the selection.
  ∘ *Warning:* Do not use “Clear Outside,” as this action is destructive to the rest of the image data.
3. Iteration: Continue this process until all significant artifacts are removed from the merged view.

##### 5. Export and Directory Structure

1. Channel Splitting: Once cleaning is complete, revert the merged image to individual channels via Image > Color > Split Channels.
2. Saving: Export each cleaned channel via File > Save As > Tiff. Adhere strictly to the naming convention established in Step 1.
3. Data Organization: Organize cleaned images into biopsy-specific folders using the following format: sub[ID]_[Timepoint]_[Side]_clean (e.g., sub8_BL_L_clean). This structure is required for the CellProfiler input module.

##### Rationale

Image processing is a necessary step towards a smooth analysis pipeline. A clean image has is free of “abnormal regions” to produce reliable and high-quality data. We remove sections where Laminin is blurry or crossed over itself, where DAPI is not clear, and overall, any section that we would not count as cells in traditional manual analysis.

#### Adaptations

- Modify contrast adjustment and/or artifact removal in accordance with immunohistochemistry methods used.
- Adjust file naming conventions in compatibility with your project. Adjustments in file naming conventions at this step will require downstream adaptations to *CellProfile*r and Python script pipeline.

## II. Cross-sectional Myofiber Object Segmentation with *CellProfiler*

### 1. Verifying pipeline with CellProfiler GUI

Before full-scale deployment, the modular pipeline must be validated within the *CellProfiler* Graphic User Interface (GUI) to ensure that segmentation parameters, specifically deep-learning thresholds and intensity-based filters, are optimized for the specific optical characteristics of the dataset.

#### Setup

1. Install **CellProfiler v4.2.8** and **Docker Desktop** from web browser. Docker is required to host the Cellpose container, ensuring a stable environment for the cyto2 and nuclei models.
2. **Plugin Configuration**: Download the runcellpose.py module from the CellProfiler-plugins repository (https://github.com/CellProfiler/CellProfiler-plugins) and move it to the local plugins directory. In *CellProfiler*, navigate to Preferences and set the **Plugins Directory** to this path.
3. **Data Input**: Load the “Cleaned Images” directory (generated in Section I) into the **Images** module.
4. **Pipeline Initialization**: Import the .cppipe file from the Dreyer Lab GitHub repository to load the pre-configured modular architecture.

#### Procedure

1. **Metadata and Channel Assignment**: In the **Metadata** module, update the Regular Expression (RegEx) to parse subject and biopsy identifiers. Use the **NamesAndTypes** module to assign tags based on naming conventions (e.g., Lam to “laminin”, DAPI to “nuclei”).
2. **Fiber Segmentation Review**: Navigate to the **RunCellpose** module assigned to the Laminin channel. Verify that the ‘cyto2’ model is selected and that the resulting outlines align precisely with the sarcolemma.
3. **Iterative Nuclear Refinement**:
  • Step through the **GaussianFilter** (sigma = 3) to confirm the smoothing of the DAPI signal.
  • Examine the **ImageMath** module to verify that the subtraction of the 17th percentile of pixel intensities effectively suppresses low-level background fluorescence and non-specific staining.
  • Inspect the final **RunCellpose** (‘nuclei’ model) output using a flow_threshold of 0.4 and cellprob_threshold of 0.0 to ensure precise myonuclear boundary adherence.
4. **Satellite Cell Identification**: In the **IdentifyPrimaryObjects** module, confirm that the **Otsu thresholding** and pixel diameter range (16–60 pixels) accurately isolate Pax7+ signals from the sarcoplasmic background.
5. **Diagnostic Visualization**: Execute **Test Mode** and use the “Window Display” to view red-line overlays on raw signals, ensuring no significant over- or under-segmentation occurs before proceeding to headless batch processing.

##### Rationale

Direct verification in the GUI is a critical quality control step that allows researchers to visually calibrate modular variables against the biological variability of the tissue. By manually inspecting the segmentation of the first few image sets, users can ensure that the deep-learning models and background subtraction logic are performing with high fidelity, minimizing the risk of false-positive nuclear assignments or inaccurate cross-sectional area (CSA) measurements during high-throughput execution.

#### Adaptations

- **Signal Intensity Variations**: If the DAPI signal is consistently dim across a cohort, the subtraction percentile in the **ImageMath** module should be reduced (e.g., to the 10th percentile) to prevent the loss of true myonuclear signals.
- **Object Size Constraints**: Adjust the pixel diameter range in the **IdentifyPrimaryObjects** module for satellite cells if the imaging magnification or species-specific cell sizes differ from the human vastus lateralis parameters provided here.
- **Alternative Detection Modes**: For images with extremely high noise or sparse signals, users may test the newest Cellpose models or alternative detection modes within the **RunCellpose** module to improve boundary detection.

### 2. Running pipeline with CellProfiler GUI

Following successful parameter verification, the high-throughput batch processing workflow is executed within the CellProfiler GUI to automate cellular segmentation and feature extraction across large datasets.

#### Setup

1. Launch Docker Desktop and *CellProfiler*. Open CellProfiler pipeline (.cpproj).
2. **Input Directory Structure**: All “Cleaned Images” must be organized into biopsy-specific folders (e.g., `sub[ID]_[Timepoint]_[Side]_clean`) to facilitate the **Metadata** module’s parsing logic.
3. **Default Output Folder**: Set the global output directory where the pipeline will programmatically generate the hierarchical folder structure for masks and morphological data.

#### Procedure

1. **Batch Initiation**: Click the **Analyze Images** button to start sequential processing. The **Groups** module will systematically organize image sets by Subject, Day, Side, and Set number based on previously extracted metadata.
2. **Sequential Segmentation Streams**:
  ∘ **Fiber Masks**: The system invokes the Cellpose ‘cyto2’ model to generate binarized sarcolemmal masks from the Laminin channel.
  **Nuclear Masks**: The pipeline executes the background-subtraction logic— subtracting the 17th percentile of pixel intensities—followed by the Cellpose ‘nuclei’ model to produce refined myonuclear masks.
  **Satellite Cell Masks**: **IdentifyPrimaryObjects** isolates Pax7+ signals using the established Otsu thresholding and size constraints (16–60 pixels).
  **MyHC1 images**: MyHC1 images are organized into desired directory location alongside output masks for downstream analysis.
3. **Data Serialization**: CellProfiler saves grayscale object masks (label matrices) and diagnostic overlays for every channel in each image set. These outputs are organized into subdirectories (e.g., laminin_masks, nuclei_masks, satcell_masks) for downstream Python analysis.

##### Rationale

Running the pipeline through the GUI provides a semi-automated balance between high-throughput speed and rigorous quality control. This environment allows the researcher to monitor segmentation progress in real-time and ensures that the standardized hierarchical directory structure is maintained, which is a prerequisite for the custom Python processing script. While the pipeline is highly automated, manual verification of the final diagnostic overlays is maintained to correct for any missed or artificial identifications, particularly in tissue regions with variable staining quality.

#### Adaptations

- **High-Volume Processing**: For datasets exceeding several hundred image sets, users should transition to the **Headless/CLI** mode (Section II-C) to reduce memory overhead associated with GUI rendering.
- **Threshold Refinement**: If the **IdentifyPrimaryObjects** module fails to capture sparse Pax7+ signals during a run, the intensity threshold can be manually overridden in the module settings to accommodate specific batch variations.
- **Alternative Identification**: This modular workflow can be easily adapted to identify other markers (e.g., macrophages) by swapping the Pax7+ **IdentifyPrimaryObjects** module with a target-specific segmentation block.

### 3. Running pipeline with CellProfiler headless

For high-throughput processing of large-scale cohorts or deployment on high-performance computing (HPC) clusters, the pipeline is executed in “headless” mode via the Command Line Interface (CLI). This method bypasses the Graphical User Interface (GUI) to minimize memory overhead and ensure strictly reproducible batch orchestration.

#### Setup

1. **Environment Configuration:** Headless execution is supported either through a **Docker container** (v28.2.2) for desktop stability or within a localized **Conda environment** for maximum performance on HPC clusters.
2. **Dependency Installation:** Create and activate a dedicated environment (Python 3.12) (Anaconda Prompt recommended) and install the core analytical suite: `conda create --name csapipeline python=3.12` `conda activate csapipeline` `conda install cellprofiler=4.2.8` `python -m pip install numpy pandas scikit-image scikit-learn openpyxl Pillow`
3. **Plugin Pathing**: Ensure the runcellpose.py plugin is located in a dedicated directory accessible to the CLI.

#### Procedure

1. **Command Invocation**: Execute the analysis using the --run (-r) and --headless (-c) flags. A standard execution string for the lab workstation follows: & “C:\Program Files\CellProfiler\CellProfiler.exe” -c -r \ `--log-level DEBUG \` `-p [PIPELINE.cppipe] \` `-i [INPUT_FOLDER] \` `-o [OUTPUT_FOLDER] \` `--plugins-directory=[ACTIVE_PLUG-INS_FOLDER]`
2. **Metadata Parsing**: The input directory path (‘-i’) must point to the root directory containing biopsy-specific subfolders. The CLI automatically applies the Regular Expression (RegEx) patterns defined in the .cpproj pipeline to organize masks and metadata.

##### Rationale

Headless execution is a mechanical necessity for high-throughput bioinformatics, as it eliminates the memory cost associated with real-time image rendering. This modality provides a superior platform for processing over 1,000 cells per biopsy, ensuring that identical parameters, such as the GaussianFilter (sigma = 3) and the 17th percentile background subtraction, are applied with absolute consistency across the entire dataset. Furthermore, native Python execution on high-performance clusters yields optimal processing speeds for multi-terabyte datasets compared to traditional desktop environments.

#### Adaptations

- **Scaling and Magnification**: While the standard pipeline is optimized for 20x magnification, researchers utilizing 10x or 40x objectives must adjust the “Estimated XY Diameter” within the RunCellpose modules to prevent over- or under-segmentation.
- **Fiber Typing Sensitivity**: The MyHC1 intensity threshold is set at a default of 25 in the downstream Python script, but this can be adjusted within the supplemental protocol if specific staining batches exhibit higher background fluorescence.
- **Marker Flexibility**: The modular design allows for the replacement of the Pax7+ identification module with other markers, such as macrophage-specific antibodies, by adjusting the pixel diameter range and thresholding strategy in the *IdentifyPrimaryObjects* module.

## III. Analysis with *Python*

### A) Run the Python Analysis

The final stage of the workflow transitions from raw segmentation masks to refined, biologically relevant datasets through a custom Python-based processing pipeline. This pipeline automates the integration of multi-channel spatial data, ensuring objective fiber-type assignment and high-throughput quantification.

#### Setup

1. **Environment Initialization:** Create and activate a dedicated environment (Python 3.12) (Anaconda Prompt recommended) and install the core analytical suite: `conda create --name csapipeline python=3.12` `conda activate csapipeline` `python -m pip install numpy pandas opencv-python XlsxWriter`
2. **Script Preparation**: Save the execution script, 23_WU_CSA_ANALYSIS.py, in your primary working directory.
3. **Input Organization**: Ensure that the outputs from CellProfiler are organized into a strict hierarchical directory structure:

~~~
    Subject/
            Timepoint/
                   L or R/
                      laminin_masks/
                      nuclei_masks/
                      satcell_masks/
                      MyHC1_images/
                      …
~~~

#### Procedure

1. Run the pipeline. From the activated Conda environment terminal, run the script with desired arguments:

~~~
python 23_WU_CSA_ANALYSIS.py <input_dir> <output_dir> <subject_name>
~~~

##### Rationale

The custom Python pipeline is a mechanical necessity to bridge the gap between computer vision outputs and biological relevance. By utilizing “greatest overlap” logic, the pipeline maintains high-fidelity assignments even in irregular or compressed fiber morphologies. Furthermore, automating fiber-type assignment via the calculate_average_intensity function removes the subjectivity inherent in manual fluorescence intensity estimation, ensuring that classification is strictly based on quantified signal distributions.

#### Adaptation

- **Scaling Factors**: Users must adjust the scaling constant within the `calculate_area_and_perimeter` function if imaging magnification differs from the 20x standard used in this study.
- **Rolling Ball Radius**: While the rolling ball radius for background subtraction is set at 100 pixels, users should adjust this value to exceed the largest cell, in cross-sectional area, present in the dataset.
- **Threshold Calibration**: While the Type I intensity threshold is set at 20, users should perform a distribution analysis (as seen in Figure 6 of the manuscript) to determine the optimal “elbow” for different staining batches.
- **Naming Conventions**: The dynamic directory parser relies on the naming convention established during pre-processing; any modifications to file nomenclature in FIJI will require corresponding updates to the RegEx patterns within the Python script (as with the CellProfiler pipeline).

### B) Output Files and Their Interpretation

The pipeline yields two primary data structures for research application and quality control (QC):

- **Subject Workbooks**: A unified Excel (.xlsx) file is generated for each subject, containing individual sheets per biopsy. Each sheet provides a morphological profile for every segmented fiber, including unique Fiber ID, Cross-Sectional Area (CSA), Perimeter, Myonuclear count, Satellite Cell count, and binary Fiber Type classification.
- **Diagnostic Overlays**: Labeled Laminin masks are saved as .tiff files to allow for rapid visual QC. These images feature unique numerical identifiers for every fiber, enabling researchers to cross-reference quantitative data with original histology.
- **Data Interpretation and Filtration**: During final analysis, users must sort and remove data labeled as “Fiber 0,” as this category represents extraneous assignments. Furthermore, any fiber assigned more than 10 nuclei must be manually verified using the diagnostic overlays to ensure segmentation accuracy.

## IV. Our Computer Specifications

This pipeline was trained and developed on both a standard, in-lab desktop workstation and a high-performance computer operated in the <u>Genomics and Cell Characterization Core Facility (G3CF) at the University of Oregon</u>.

### Lab desktop

Dell Precision 3630 Tower workstation running Microsoft Windows 11 Enterprise (Build 26100). The system is equipped with an Intel® Core™ i7-9700 CPU @ 3.00 GHz with 8 cores and 8 threads and operates in UEFI mode with Secure Boot enabled. The machine has a 64-bit architecture and BIOS version 2.26.0 (dated December 8, 2023).

### G3CF computer

32-core AMD Ryzen Threadripper 3970X processor running at 3.9 GHz and 256 GB of DDR4 RAM. The system features a custom water-cooled configuration optimized for high-throughput image processing and is powered by an NVIDIA GeForce RTX 5090 graphics card to support GPU-accelerated rendering and analysis tasks. The machine runs Microsoft Windows 10 Enterprise (version 10.0.19045, build 19045) in UEFI BIOS mode. The software environment includes Imaris 9.9 with the Cell and Filaments Modules Imaris 9.9 Converter, MATLAB 2018b, and Zeiss Zen Black.

## V. *CellProfiler* Pipeline Module Parameters

*Pipeline: 23_WU_CSA_PIPELINE.cpproj* | *CellProfiler 4.2.8*

| Step | Module | Parameter | Value |
| --- | --- | --- | --- |
| #01 | **RunCellpose**<br>*Cellpose for laminin fibers* | Rescale images before running Cellpose | Yes ● |
|  |  | Run CellPose in docker or local python environment | Docker |
|  |  | Select Cellpose docker image | cellprofiler/runcellpose_with_pretrained:2.3.2 |
|  |  | Detection mode | cyto2 |
|  |  | Supply nuclei image as well? | No ● |
|  |  | Select the input image | laminin |
|  |  | Expected object diameter | 180 |
|  |  | Cell probability threshold | 0.0 |
|  |  | Minimum size | 40 |
|  |  | Flow threshold | 0.4 |
|  |  | Name the output object | laminin_identified |
|  |  | Invert images | No ● |
|  |  | Save probability image? | No ● |
|  |  | Use 3D | No ● |
|  |  | Stitch Threshold | 0.0 |
| #02 | **ConvertObjectsToImage** | Select the input objects | laminin_identified |
|  |  | Name the output image | laminin_identified_mask |
|  |  | Select the color format | Grayscale |
| #03 | **SaveImages** | Select the type of image to save | Image |
|  |  | Select the image to save | laminin_identified_mask |
|  |  | Select method for constructing file names | From image filename |
|  |  | Select image name for file prefix | laminin (from NamesAndTypes) |
|  |  | Append a suffix to the image file name? | Yes ● |
|  |  | Text to append to the image name | _laminin_masks |
|  |  | Saved file format | tiff |
|  |  | Image bit depth | 8-bit integer |
|  |  | Save with lossless compression? | Yes ● |
|  |  | Output file location | Default Output Folder sub-folder |
|  |  | Sub-folder | ./Subject/Day/Side/laminin_masks |
|  |  | Overwrite existing files without warning? | No ● |
|  |  | When to save | Every cycle |
|  |  | Record the file and path information to the saved image? | No ● |
|  |  | Create subfolders in the output folder? | No ● |
| #04 | **OverlayOutlines**<br>*Overlays Fibers identified with laminin* | Display outlines on a blank image? | No ● |
|  |  | Select image on which to display outlines | laminin (from NamesAndTypes) |
|  |  | Name the output image | laminin_fiber_image |
|  |  | Outline display mode | Color |
|  |  | How to outline | Inner |
|  |  | Select objects to display | laminin_identified (from RunCellpose) |
|  |  | Select outline color | Red |
| #05 | **SaveImages**<br>*Saves laminin and identified fiber overlay* | Select the type of image to save | Image |
|  |  | Select the image to save | laminin_fiber_image (from OverlayOutlines) |
|  |  | Select method for constructing file names | From image filename |
|  |  | Select image name for file prefix | laminin (from NamesAndTypes) |
|  |  | Append a suffix to the image file name? | Yes ● |
|  |  | Text to append to the image name | _laminin_fibers |
|  |  | Saved file format | tiff |
|  |  | Image bit depth | 8-bit integer |
|  |  | Save with lossless compression? | Yes ● |
|  |  | Output file location | Default Output Folder sub-folder |
|  |  | Sub-folder | ./Subject/Day/Side/laminin_overlay |
|  |  | Overwrite existing files without warning? | No ● |
|  |  | When to save | Every cycle |
|  |  | Record the file and path information to the saved image? | No ● |
|  |  | Create subfolders in the output folder? | No ● |
| #06 | **GaussianFilter**<br>*GaussianFilter on nuclei image* | Select the input image | nuclei (from NamesAndTypes) |
|  |  | Name the output image | gaussian_nuclei |
|  |  | Sigma | 3 |
| #07 | **RunCellpose**<br>*Identified nuclei in the DAPI image* | Rescale images before running Cellpose | Yes ● |
|  |  | Run CellPose in docker or local python environment | Docker |
|  |  | Select Cellpose docker image | cellprofiler/runcellpose_with_pretrained:2.3.2 |
|  |  | Detection mode | nuclei |
|  |  | Select the input image | gaussian_nuclei (from GaussianFilter) |
|  |  | Expected object diameter | 15 |
|  |  | Cell probability threshold | 0.0 |
|  |  | Minimum size | 20 |
|  |  | Flow threshold | 0.4 |
|  |  | Name the output object | cellpose_nuclei |
|  |  | Invert images | No ● |
|  |  | Save probability image? | No ● |
|  |  | Use 3D | No ● |
|  |  | Stitch Threshold | 0.0 |
| #08 | **MeasureImageIntensity** | Select images to measure | gaussian_nuclei |
|  |  | Measure the intensity only from areas enclosed by objects? | Yes ● |
|  |  | Select input object sets | cellpose_nuclei |
|  |  | Calculate custom percentiles | Yes ● |
|  |  | Specify percentiles to measure | 15, 16, 17 |
| #09 | **ImageMath**<br>*Subtract 12th percentile of intensity from gaussian image* | Operation | Subtract |
|  |  | Name the output image | _subtracted_gaussian_nuclei |
|  |  | Image or measurement? (1st operand) | Image |
|  |  | Select the first image | gaussian_nuclei (from GaussianFilter) |
|  |  | Multiply the first image by | 1.0 |
|  |  | Image or measurement? (2nd operand) | Measurement |
|  |  | Category | Intensity |
|  |  | Measurement: Measurement | Percentile_17 |
|  |  | Measurement: Image | gaussian_nuclei_cellpose_nuclei |
|  |  | Multiply the second image by | 1.0 |
|  |  | Raise the power of the result by | 1.0 |
|  |  | Multiply the result by | 1.0 |
|  |  | Add to result | 0.0 |
|  |  | Set values less than 0 equal to 0? | Yes ● |
|  |  | Set values greater than 1 equal to 1? | Yes ● |
|  |  | Replace invalid values with 0? | Yes ● |
| #10 | **RunCellpose**<br>*Rerun Cellpose on subtracted Gaussian image* | Rescale images before running Cellpose | Yes ● |
|  |  | Run Cellpose in docker or local python environment | Docker |
|  |  | Select Cellpose docker image | cellprofiler/runcellpose_with_pretrained:2.3.2 |
|  |  | Detection mode | nuclei |
|  |  | Select the input image | _subtracted_gaussian_nuclei (from ImageMath) |
|  |  | Expected object diameter | 25 |
|  |  | Cell probability threshold | 0.0 |
|  |  | Minimum size | 30 |
|  |  | Flow threshold | 0.4 |
|  |  | Name the output object | cellpose_subtracted_nuclei |
|  |  | Invert images | No ● |
|  |  | Save probability image? | No ● |
|  |  | Use 3D | No ● |
|  |  | Stitch Threshold | 0.0 |
| #11 | **ConvertObjectsToImage** | Select the input objects | cellpose_subtracted_nuclei |
|  |  | Name the output image | filtered_nuclei |
|  |  | Choose the color format | Grayscale |
| #12 | **SaveImages**<br>*Save filtered_nuclei image* | Select the type of image to save | Image |
|  |  | Select the image to save | filtered_nuclei (from ConvertObjectsToImage) |
|  |  | Select method for constructing file names | From image filename |
|  |  | Select image name for file prefix | nuclei (from NamesAndTypes) |
|  |  | Append a suffix to the image file name? | Yes ● |
|  |  | Text to append to the image name | _filtered_nuclei |
|  |  | Saved file format | tiff |
|  |  | Image bit depth | 8-bit integer |
|  |  | Save with lossless compression? | Yes ● |
|  |  | Output file location | Default Output Folder sub-folder |
|  |  | Sub-folder | ./Subject/Day/Side/filtered_nuclei |
|  |  | Overwrite existing files without warning? | Yes ● |
|  |  | When to save | Every cycle |
|  |  | Record the file and path information to the saved image? | No ● |
|  |  | Create subfolders in the output folder? | No ● |
| #13 | **ConvertObjectsToImage**<br>*Convert nuclei objects to image (nuclei_masks)* | Select the input objects | cellpose_subtracted_nuclei |
|  |  | Name the output image | nuclei_masks |
|  |  | Select the color format | Grayscale |
| #14 | **SaveImages**<br>*Save nuclei_masks image* | Select the type of image to save | Image |
|  |  | Select the image to save | nuclei_masks (from ConvertObjectsToImage) |
|  |  | Select method for constructing file names | From image filename |
|  |  | Select image name for file prefix | nuclei (from NamesAndTypes) |
|  |  | Append a suffix to the image file name? | Yes ● |
|  |  | Text to append to the image name | _nuclei_masks |
|  |  | Saved file format | tiff |
|  |  | Image bit depth | 8-bit integer |
|  |  | Save with lossless compression? | Yes ● |
|  |  | Output file location | Default Output Folder sub-folder |
|  |  | Sub-folder | ./Subject/Day/Side/nuclei_masks |
|  |  | Overwrite existing files without warning? | No ● |
|  |  | When to save | Every cycle |
|  |  | Record the file and path information to the saved image? | No ● |
|  |  | Create subfolders in the output folder? | No ● |
| #15 | **OverlayOutlines**<br>*cleaned/raw DAPI and nuclei identified overlay* | Display outlines on a blank image? | No ● |
|  |  | Select image on which to display outlines | nuclei |
|  |  | Name the output image | nuclei_over_dapi |
|  |  | Outline display mode | Color |
|  |  | How to outline | Inner |
|  |  | Select objects to display | cellpose_subtracted_nuclei |
|  |  | Select outline color | Red |
| #16 | **SaveImages**<br>*Cleaned/raw DAPI and nuclei identified overlay* | Select the type of image to save | Image |
|  |  | Select the image to save | nuclei_over_dapi (from OverlayOutlines) |
|  |  | Select method for constructing file names | From image filename |
|  |  | Select image name for file prefix | nuclei (from NamesAndTypes) |
|  |  | Append a suffix to the image file name? | Yes ● |
|  |  | Text to append to the image name | nuclei_over_dapi |
|  |  | Saved file format | tiff |
|  |  | Image bit depth | 8-bit integer |
|  |  | Save with lossless compression? | Yes ● |
|  |  | Output file location | Default Output Folder sub-folder |
|  |  | Sub-folder | ./Subject/Day/Side/nuclei_overlay |
|  |  | Overwrite existing files without warning? | No ● |
|  |  | When to save | Every cycle |
|  |  | Record the file and path information to the saved image? | No ● |
|  |  | Create subfolders in the output folder? | No ● |
| #17 | **GaussianFilter**<br>*GaussianFilter on satellite image* | Select the input image | satellite (from NamesAndTypes) |
|  |  | Name the output image | gaussian_satellite |
|  |  | Sigma | 2 |
| #18 | **IdentifyPrimaryObjects** | Use advanced settings? | Yes ● |
|  |  | Select the input image | gaussian_satellite (from GaussianFilter) |
|  |  | Name the primary objects to be identified | satcells |
|  |  | Typical diameter of objects, in pixel units (Min, Max) | 16, 60 |
|  |  | Discard objects outside the diameter range? | No ● |
|  |  | Discard objects touching the border of the image? | Yes ● |
|  |  | Threshold strategy | Global |
|  |  | Thresholding method | Otsu |
|  |  | Two-class or three-class thresholding? | Three classes |
|  |  | Assign pixels in the middle intensity class to the foreground or the background? | Background |
|  |  | Threshold smoothing scale | 0 |
|  |  | Threshold correction factor | 1.0 |
|  |  | Lower and upper bounds on threshold | 0.24 1.0 |
|  |  | Log transform before thresholding? | No ● |
|  |  | Method to distinguish clumped objects | Intensity |
|  |  | Method to draw dividing lines between clumped objects | Intensity |
|  |  | Automatically calculate size of smoothing filter for declumping? | Yes ● |
|  |  | Automatically calculate minimum allowed distance between local maxima? | Yes ● |
|  |  | Speed up by using lower-resolution image to find local maxima? | No ● |
|  |  | Display accepted local maxima? | No ● |
|  |  | Fill holes in identified objects? | After both thresholding and declumping |
| #19 | **OverlayOutlines**<br>*Satellite cell overlay on satellite image* | Display outlines on a blank image? | No ● |
|  |  | Select image on which to display outlines | satellite (from NamesAndTypes) |
|  |  | Name the output image | satcell_overlay |
|  |  | Outline display mode | Color |
|  |  | How to outline | Inner |
|  |  | Select objects to display | satcells (from IdentifyPrimaryObjects) |
|  |  | Select outline color | Red |
| #20 | **SaveImages**<br>*Save satcell_overlay* | Select the type of image to save | Image |
|  |  | Select the image to save | satcell_overlay (from OverlayOutlines) |
|  |  | Select method for constructing file names | From image filename |
|  |  | Select image name for file prefix | satellite (from NamesAndTypes) |
|  |  | Append a suffix to the image file name? | No ● |
|  |  | Text to append to the image name | (empty) |
|  |  | Saved file format | tiff |
|  |  | Image bit depth | 8-bit integer |
|  |  | Save with lossless compression? | Yes ● |
|  |  | Output file location | Default Output Folder sub-folder |
|  |  | Sub-folder | ./Subject/Day/Side/satcell_outlines |
|  |  | Overwrite existing files without warning? | Yes ● |
|  |  | When to save | Every cycle |
|  |  | Record the file and path information to the saved image? | No ● |
|  |  | Create subfolders in the output folder? | No ● |
| #21 | **ConvertObjectsToImage**<br>*Convert satcells to satcell_masks (Grayscale)* | Select the input objects | satcells from IdentifyPrimaryObjects) |
|  |  | Name the output image | satcell_masks |
|  |  | Select the color format | Grayscale |
| #22 | **SaveImages**<br>*Save satcell_masks image* | Select the type of image to save | Image |
|  |  | Select the image to save | satcell_masks (from ConvertObjectsToImage) |
|  |  | Select method for constructing file names | From image filename |
|  |  | Select image name for file prefix | satellite (from NamesAndTypes) |
|  |  | Append a suffix to the image file name? | Yes ● |
|  |  | Text to append to the image name | _satcell_masks |
|  |  | Saved file format | tiff |
|  |  | Image bit depth | 8-bit integer |
|  |  | Save with lossless compression? | Yes ● |
|  |  | Output file location | Default Output Folder sub-folder |
|  |  | Sub-folder | ./Subject/Day/Side/satcell_masks |
|  |  | Overwrite existing files without warning? | Yes ● |
|  |  | When to save | Every cycle |
|  |  | Record the file and path information to the saved image? | No ● |
|  |  | Create subfolders in the output folder? | No ● |
| #23 | **SaveImages**<br>*Save MyHC1 images* | Select the type of image to save | Image |
|  |  | Select the image to save | MyHC1 (from NamesAndTypes) |
|  |  | Select method for constructing file names | From image filename |
|  |  | Select image name for file prefix | MyHC1 (from NamesAndTypes) |
|  |  | Append a suffix to the image file name? | Yes ● |
|  |  | Text to append to the image name | (empty — no suffix text entered) |
|  |  | Saved file format | tiff |
|  |  | Image bit depth | 8-bit integer |
|  |  | Save with lossless compression? | Yes ● |
|  |  | Output file location | Default Output Folder sub-folder |
|  |  | Sub-folder | ./Subject/Day/Side/MyHC1_images |
|  |  | Overwrite existing files without warning? | Yes ● |
|  |  | When to save | Every cycle |
|  |  | Record the file and path information to the saved image? | No ● |
|  |  | Create subfolders in the output folder? | No ● |

## Bibliography

1. Mazzarini M, Falchi M, Bani D, Migliaccio AR. Evolution and new frontiers of histology in bio-medical research. Microsc Res Tech. 2021;84(2):217–237. doi:10.1002/jemt.23579

2. Cheek DB, Holt AB, Hill DE, Talbert JL. Skeletal Muscle Cell Mass and Growth: The Concept of the Deoxyribonucleic Acid Unit. Pediatr Res. 1971;5(7):312–328. doi:10.1203/00006450-197107000-00004

3. Cramer AAW, Prasad V, Eftestøl E, et al. Nuclear numbers in syncytial muscle fibers promote size but limit the development of larger myonuclear domains. Nat Commun. 2020;11(1):6287. doi:10.1038/s41467-020-20058-7

4. Hansson KA, Eftestøl E, Bruusgaard JC, et al. Myonuclear content regulates cell size with similar scaling properties in mice and humans. Nat Commun. 2020;11(1):6288. doi:10.1038/s41467-020-20057-8

5. Moesgaard L, Jessen S, Mackey AL, Hostrup M. Myonuclear addition is associated with sex-specific fiber hypertrophy and occurs in relation to fiber perimeter not cross-sectional area. J Appl Physiol Bethesda Md 1985. 2022;133(3):732–741. doi:10.1152/japplphysiol.00235.2022

6. Gundersen, K. (2016). Muscle memory and a new cellular model for muscle atrophy and hypertrophy. The Journal of Experimental Biology, 219(Pt 2), 235–242. 10.1242/jeb.124495

7. Egner, I. M., Bruusgaard, J. C., Eftestøl, E., & Gundersen, K. (2013). A cellular memory mechanism aids overload hypertrophy in muscle long after an episodic exposure to anabolic steroids. The Journal of Physiology, 591(24), 6221–6230. 10.1113/jphysiol.2013.264457

8. Muyskens, J. B., Foote, D. M., Bigot, N. J., Strycker, L. A., Smolkowski, K., Kirkpatrick, T. K., Lantz, B. A., Shah, S. N., Mohler, C. G., Jewett, B. A., Owen, E. C., & Dreyer, H. C. (2019). Cellular and morphological changes with EAA supplementation before and after total knee arthroplasty. Journal of Applied Physiology (Bethesda, Md.: 1985), 127(2), 531–545. 10.1152/japplphysiol.00869.2018

9. Mackey AL, Kjaer M, Charifi N, Henriksson J, Bojsen-Moller J, Holm L, Kadi F. Assessment of satellite cell number and activity status in human skeletal muscle biopsies. Muscle Nerve. 2009 Sep;40(3):455–65. doi: 10.1002/mus.21369. PMID: 19705426.

10. Wen Y, Murach KA, Vechetti IJ Jr, Fry CS, Vickery C, Peterson CA, McCarthy JJ, Campbell KS. MyoVision: software for automated high-content analysis of skeletal muscle immunohistochemistry. J Appl Physiol (1985). 2018 Jan 1;124(1):40–51. doi: 10.1152/japplphysiol.00762.2017. Epub 2017 Oct 5. PMID: 28982947; PMCID: PMC6048460.

11. Sanz G, Martínez-Aranda LM, Tesch PA, Fernandez-Gonzalo R, Lundberg TR. Muscle2View, a CellProfiler pipeline for detection of the capillary-to-muscle fiber interface and high-content quantification of fiber type-specific histology. J Appl Physiol (1985). 2019 Dec 1;127(6):1698–1709. doi: 10.1152/japplphysiol.00257.2019. Epub 2019 Nov 7. PMID: 31697593.

12. Encarnacion-Rivera L, Foltz S, Hartzell HC, Choo H. Myosoft: An automated muscle histology analysis tool using machine learning algorithm utilizing FIJI/ImageJ software. PLoS One. 2020 Mar 4;15(3):e0229041. doi: 10.1371/journal.pone.0229041. PMID: 32130242; PMCID: PMC7055860.

13. Lau YS, Xu L, Gao Y, Han R. Automated muscle histopathology analysis using CellProfiler. Skelet Muscle. 2018 Oct 18;8(1):32. doi: 10.1186/s13395-018-0178-6. PMID: 30336774; PMCID: PMC6193305.

14. Babcock LW, Hanna AD, Agha NH, Hamilton SL. MyoSight-semi-automated image analysis of skeletal muscle cross sections. Skelet Muscle. 2020 Nov 16;10(1):33. doi: 10.1186/s13395-020-00250-5. PMID: 33198807; PMCID: PMC7667765.

15. Mayeuf-Louchart A, Hardy D, Thorel Q, Roux P, Gueniot L, Briand D, Mazeraud A, Bouglé A, Shorte SL, Staels B, Chrétien F, Duez H, Danckaert A. MuscleJ: a high-content analysis method to study skeletal muscle with a new Fiji tool. Skelet Muscle. 2018 Aug 6;8(1):25. doi: 10.1186/s13395-018-0171-0. PMID: 30081940; PMCID: PMC6091189

